# Mechanistic Modeling Explains the Production Dynamics of Recombinant Adeno-Associated Virus with the Baculovirus Expression Vector System

**DOI:** 10.1101/2023.02.04.527082

**Authors:** Francesco Destro, Prasanna Srinivasan, Joshua M. Kanter, Caleb Neufeld, Jacqueline M. Wolfrum, Paul W. Barone, Stacy L. Springs, Anthony J. Sinskey, Sylvain Cecchini, Robert M. Kotin, Richard D. Braatz

**Affiliations:** Department of Chemical Engineering, Massachusetts Institute of Technology, Cambridge, MA; Center for Biomedical Innovation, Massachusetts Institute of Technology, Cambridge, MA; Gene Therapy Center, University of Massachusetts Chan Medical School, Worcester, MA; Department of Biology, Massachusetts Institute of Technology, Cambridge, MA; Carbon Biosciences, Waltham, MA

**Keywords:** gene therapy, viral vector, mechanistic modeling, biomanufacturing, baculovirus expression vector system, AAV-based gene therapy, recombinant adeno-associated virus, viral vector manufacturing

## Abstract

The demand for recombinant adeno-associated virus (rAAV) for gene therapy is expected to soon exceed current manufacturing capabilities, considering the expanding number of approved products and of pre-clinical and clinical stage studies. Current rAAV manufacturing processes have less-than-desired yields and produce a significant amount of empty capsids. Recently, FDA approved the first rAAV-based gene therapy product manufactured in the baculovirus expression vector system (BEVS). The BEVS technology, based on an invertebrate cell line derived from *Spodoptera frugiperda*, demonstrated scalable production of high volumetric titers of full capsids. In this work, we develop a mechanistic model describing the key extracellular and intracellular phenomena occurring during baculovirus infection and rAAV virion maturation in the BEVS. The predictions of the model show good agreement with experimental measurements reported in the literature on rAAV manufacturing in the BEVS, including for TwoBac, ThreeBac, and OneBac constructs. The model is successfully validated against measured concentrations of structural and non-structural protein components, and of vector genome. We carry out a model-based analysis of the process, to provide insights on potential bottlenecks that limit the formation of full capsids. The analysis suggests that vector genome amplification is the limiting step for rAAV production in TwoBac. In turn, vector genome amplification is limited by low Rep78 levels. For ThreeBac, low vector genome amplification dictated by Rep78 limitation appears even more severe than in TwoBac. Transgene expression in the insect cell during rAAV manufacturing is also found to negatively influence the final rAAV production yields.

## Introduction

Recombinant adeno-associated virus (rAAV) is one of the most important vectors for in vivo and ex vivo gene therapy. The first gene therapy treatments approved in the U.S.^1^ and in the E.U.^2^ rely on rAAV vectors, and currently more than 200 clinical trials worldwide involve rAAV-based therapies.^3^ Considering that up to 30 new rAAV therapies are expected to be launched by 2025,^4^ the demand for rAAV is estimated to soon exceed the production capacity, potentially limiting access to these therapeutics for both pivotal pre-clinical and clinical trials and for treating patients in need.^5^ This issue motivates developing more efficient processes for rAAV manufacturing.

State-of-the-art technology for rAAV manufacturing is based on either mammalian or insect cell lines, to which the genes for AAV production are introduced through transfection or infection processes. The most productive processes for mammalian cells are based on suspension cultures with AAV genes introduced using either transient transfection with plasmids or infection with recombinant human herpes simplex virus type 1.^6,7^ These systems also produce large amounts of empty rAAV capsids (up to 80–95% of the total),^8,9^ which are associated with a low rAAV volumetric titer and the extra cost and processes for enriching the filled particles before delivery to the patient. In contrast, rAAV manufacturing through recombinant baculovirus (BV) infection of insect cell lines can reliably achieve filled-to-empty capsid ratios of 50–80%.^10,11^ The cell line derived from *Spodoptera frugiperda* (Sf9) is conventionally used with the baculovirus expression vector system (BEVS).^12,13^ Compared to mammalian cell-based processes, insect cells have multiple advantages. Production of recombinant proteins with the BEVS in Sf9 suspension cultures is well established, including for vaccine manufacturing.^14^ In addition, insect cells are intrinsically more resistant to contamination from human pathogens than mammalian cell lines,^15^ and, in insect cells, many mammalian promoters are inactive or attenuated, hence detrimental effects caused by transgene expression during rAAV production are averted. Baculovirus infection propagates through virus budding in the same cells that produce rAAV, resulting in near quantitative infection of the entire cell culture. In contrast, transient transfection does not result in cell-to-cell spread of plasmid, hence transfection efficiency is a major determinant of productivity. Multiple clinical rAAV vectors for gene therapies are already produced in Sf9 cells with the BEVS,^2^ and FDA recently approved for the first time a gene therapy product manufactured with the BEVS (HEMGENIX^®^).^16,17^

Different BEVS systems have been developed to produce rAAV with Sf9 cells through infection from recombinant *Autographa californica* multiple nuclear polyhedrosis viruses (AcMNPV): ThreeBac,^18^ TwoBac,^19^ OneBac^20^ and MonoBac^21^. In each of the systems, the genetic elements for producing rAAV (*Rep, Cap* and the vector genome template)^7^ are split amongst different recombinant baculoviruses, or integrated into the host genome. The first study demonstrating production of rAAV in insect cells introduced the ThreeBac system,^18^ involving coinfection of Sf9 cells with three viruses (Fig. S1). The three recombinant baculoviruses (repBV, capBV, and goiBV) carry the genetic information necessary for rAAV production^7^ in four cassettes (Fig. 1a): the *Rep52* and *Rep78* cassettes (in repBV), the *Cap* cassette (in capBV), and the inverted terminal repeat/gene of interest (ITR/GOI) cassette (in goiBV). The ThreeBac system achieved rAAV production per cell comparable to the production achieved with mammalian cell processes, but has the advantage that Sf9 cells have the potential to grow to higher cell density.^9^ The ThreeBac system demonstrated poor passage stability, due to tandem duplication of homologous regions of the *Rep52* and *Rep78* genes in repBV.

**Figure 1:**
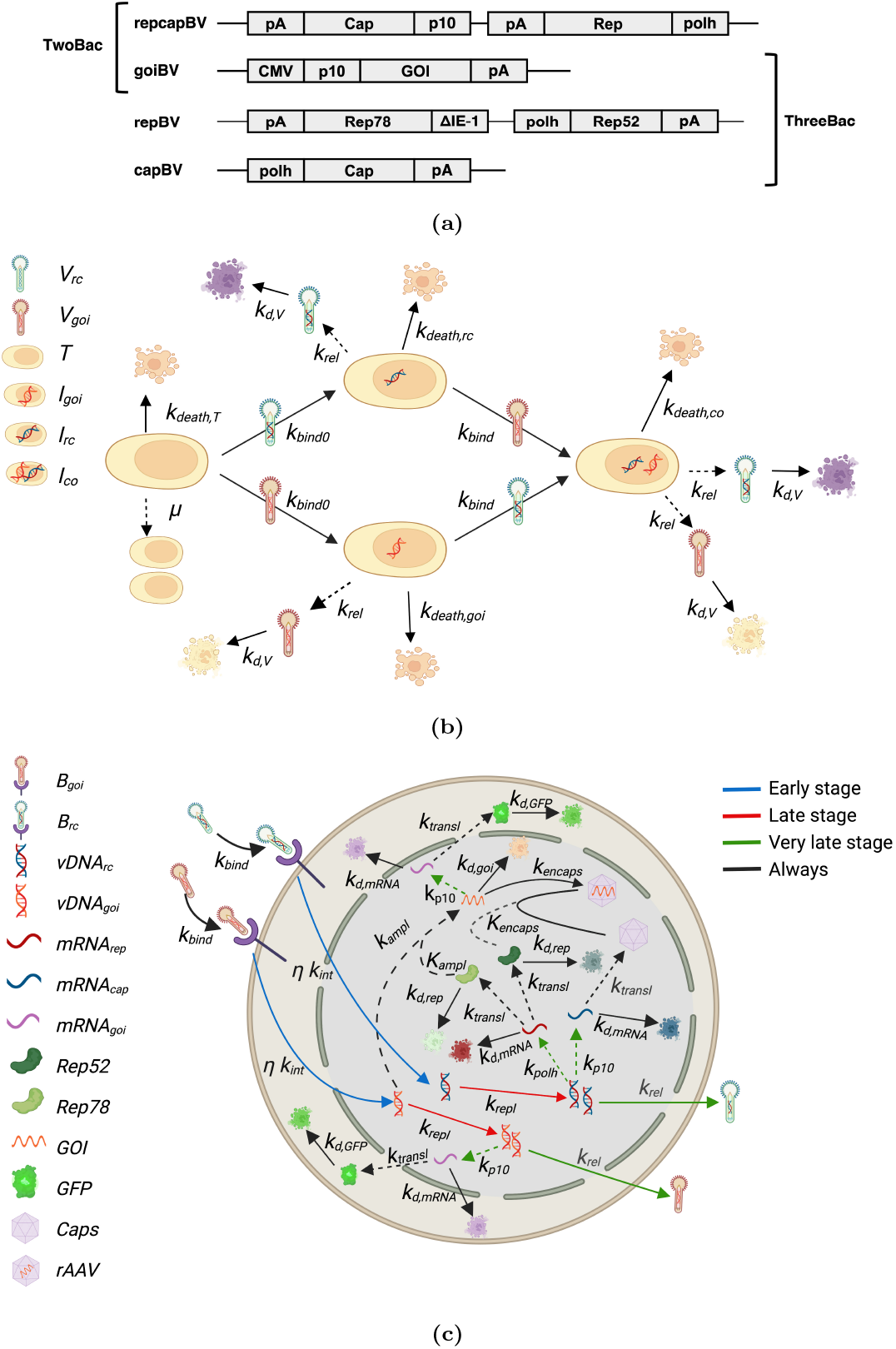
(a) TwoBac and ThreeBac constructs. TwoBac delivers the genetic information for rAAV production through two baculoviruses: goiBV and repcapBV. The goiBV carries the ITR/GOI cassette, while the repcapBV contains a *Rep* cassette, from which both Rep52 and Rep78 are expressed, and a *Cap* cassette, from which the structural proteins are expressed. In ThreeBac, the same goiBV as in TwoBac is used, but *Rep* and *Cap* are delivered through two separate baculoviruses: repBv and capBV, respectively. In repBV, Rep proteins are expressed through separate *Rep52* and *Rep78*. The promoters in all cassettes in TwoBac and ThreeBac are the strong *Autographa californica* very late promoters polh or p10, except for the *Rep78* cassette in repBV, which contains the *Orgya pseudotsugata* weak immediate early promoter ΔIE1 instead. (b) TwoBac: baculovirus infection model. Uninfected cells duplicate with kinetic constant *μ*, and present a low death kinetics (*k*_*death,T*_). Virions in the medium bind uninfected cells with kinetic constant *k*_*bind*0_. Infected cells are infected by additional virus with decreased binding kinetics (*k*_*bind*_). Coinfection from both repcapBV and goiBV is necessary for rAAV production. Infected cells do not duplicate, and experience an accelerated death kinetics. Budded virions are released in the very late infection stage with rate *k*_*rel*_. Virions in the medium slowly degrade (*k*_*d,V*_). (c) TwoBac: intracellular reaction-transport network. Receptor-bound baculovirus is transported into the nucleus (*k*_*int*_). Rerouting to lysosomes leads to degradation of several (*η*) baculoviruses. In the late stage of the infection, the viral DNA replicates (*k*_*repl*_) in the nucleus. Transcription from the promoters used in TwoBac (polh and p10) occurs during the very late infection stage. The two promoters present slightly different transcription kinetics (*k*_*p*10_ and *k*_*polh*_). Rep52 and Rep78 are synthetized (*k*_*transl*_) from the *Rep* transcript, while empty rAAV capsids form from the structural proteins translated from the *Cap* transcript. The transgene is expressed through the p10 promoter. Vector genome amplification (*k*_*ampl*_) originates from the ITR/GOI cassette of goiBV. Rep78 and Rep52 play a role in, respectively, vector genome amplification (*K*_*ampl*_) and encapsidation (*k*_*encaps*_). Non-negligible degradation is registered for Rep proteins (*k*_*d,rep*_), for mRNA (*k*_*d,mRNA*_), and for the non-encapsidated vector genome (*k*_*d,DNA*_). Dashed lines indicate that reactants are not consumed in the reaction. Figures 1b and 1c were created with BioRender (https://biorender.com/).

Since then, different improved BEVSs for rAAV manufacturing have been proposed.^22,23^ The TwoBac system^19^ was the first set of constructs to overcome the passage-instability limitation of ThreeBac. In TwoBac, Sf9 cells are coinfected by only two recombinant baculoviruses: repcapBV and goiBV (Fig. 1). While goiBV in TwoBac is equivalent to the one used in ThreeBac, repcapBV features both *Rep* and *Cap* open reading frames (ORFs). The *Rep* cassette is designed to express both *Rep78* and *Rep52* (in approximately equivalent levels) from a single transcript, while the *Cap* ORF, again from a single transcript, expresses AAV structural proteins (VP1, VP2, and VP3) in appropriate stoichiometric ratios to mimic the capsid composition of wild-type AAV. Recently, rAAV production with BEVSs using only one recombinant baculovirus has been demonstrated. In the OneBac system,^20^ an insect cell line with stably integrated *Rep* and *Cap* cassettes is infected by goiBV. In the MonoBac system,^21^ instead, Sf9 cells are infected by a recombinant baculovirus carrying all the cassettes necessary for rAAV production.

Although rAAV manufacturing with the BEVS is a promising technology, many challenges remain to be addressed. Mechanistic modeling is an invaluable tool to support (bio)pharmaceutical process development,^24,25^ as it allows a faster understanding of the process dynamics with less time-consuming and expensive experiments, compared to traditional process development. Recently, mechanistic modeling of rAAV production via transient transfection of mammalian cells identified bottlenecks in the process, and was used to propose improvements to the system.^26^ Mathematical models for baculovirus infection and production of recombinant proteins and virus-like particles have been proposed in the past.^27–35^ However, to the best of our knowledge, no mathematical model has been developed to describe and improve rAAV production with the BEVS, a process much more complex than standard manufacturing of recombinant proteins.

In this work, we develop a mechanistic model for baculovirus infection and rAAV production from insect cells. The model is developed for two-wave synchronous infection, namely it can simulate the process when baculovirus multiplicity of infection (MOI) is large enough to infect all cells of the batch shortly after that infectious baculovirus is added to the system (first wave), or, at most, after that budded baculovirus is released from the first wave of infected cells (second wave). The model considers the intermediate steps of baculovirus binding, baculovirus transport to the nucleus and replication, release of budded baculovirus, transcription and translation of AAV genes, rAAV capsid formation, Rep protein synthesis, vector genome amplification, and vector genome encapsidation. The parameters of the model are estimated from data available in the literature. The model is extensively validated against experimental datasets from the literature that report dynamic measurements of intermediates and rAAV titer collected during batch processing for TwoBac and ThreeBac. The model is also validated against literature data collected from stably transfected cell lines. An in-silico process analysis is carried out, including model simulation, sensitivity analysis, and uncertainty analysis, to investigate the productivity bottlenecks for TwoBac and ThreeBac, namely the systems for which the model has been more extensively validated. Potential strategies are discussed for increasing productivity for TwoBac, currently the most widely used BEVS construct. Finally, a differential analysis is carried out between modeling rAAV production in insect and mammalian cell lines.

## Results

A model is developed for the batch production of rAAV in Sf9 cells using the BEVS. Taking as reference the TwoBac process, the formulation of the model is outlined in the Mathematical Modeling Formulation section in Materials and Methods, which also summarizes the differences across TwoBac, ThreeBac, and stable cell lines production modeling. The model features 32 parameters (Table 1), which are either retrieved from the literature or estimated from experimental data, as outlined in Materials and Methods. The TwoBac and ThreeBac constructs are shown in Fig. 1a. The steps and species considered by the model for TwoBac and ThreeBac are shown in Figs. 1 and S1, respectively. In the remainder of this manuscript, we interchangeably refer to full rAAV capsids as filled capsids or encapsidated vector genomes, as opposed to empty capsids.

**Table 1:** Model parameters

| Symbol | Value | 95% confidence interval | Unit | Parameter | Source |
| --- | --- | --- | --- | --- | --- |
| <u>Cell growth and death dynamics</u> |  |  |  |  |  |
| $k_{death,cap}$ | 0 | – | $h^{-1}$ | Infected cell death kinetics: dependence on rAAV capsids | Estimated from literature data<sup>[44]</sup> |
| $k_{death,DNA}$ | $2.9 \times 10^{-3}$ | $(2.5 \times 10^{-3}, 3.3 \times 10^{-3})$ | $h^{-1}$ | Infected cell death kinetics: dependence on baculovirus DNA | Estimated from literature data<sup>[44]</sup> |
| $k_{death,rep}$ | 0 | – | $h^{-1}$ | Infected cell death kinetic constant: dependence on Rep78 | Estimated from literature data<sup>[44]</sup> |
| $k_{death,T}$ | $8 \times 10^{-5}$ | $(6 \times 10^{-5}, 1 \times 10^{-4})$ | $h^{-1}$ | Uninfected cell death kinetic constant | From literature<sup>[30]</sup> |
| $\mu$ | $2.8 \times 10^{-2}$ | $(2.7 \times 10^{-2}, 2.9 \times 10^{-2})$ | $h^{-1}$ | Uninfected cell growth kinetic constant | From literature<sup>[30,46]</sup> |
| $\tau_{death}$ | 24 | (22, 26) | hpi | Switch from uninfected to infected death kinetics | Estimated from literature data<sup>[44]</sup> |
| <u>Binding and transport to nucleus</u> |  |  |  |  |  |
| $k_{bind0}$ | $6.3 \times 10^{-7}$ | $(5.2 \times 10^{-7}, 7.4 \times 10^{-7})$ | $mL \text{ cell}^{-1} h^{-1}$ | Uninfected cell binding kinetic constant | Estimated from literature data<sup>[31]</sup> |
| $k_{int}$ | 0.60 | (0.48, 0.72) | $h^{-1}$ | Baculovirus internalization kinetic constant | From literature<sup>[28]</sup> |
| $\beta_{bind}$ | 0.149 | (0.120, 0.178) | $h^{-1}$ | Infected cell binding decay kinetic constant | Estimated from literature data<sup>[31]</sup> |
| $\eta$ | 0.50 | (0.45, 0.55) | – | Percentage of baculovirus rerouted to lysosomes | From literature<sup>[28]</sup> |
| $\tau_{bind}$ | 1.8 | (1.3, 2.3) | $h^{-1}$ | Infected cell binding decay onset | Estimated from literature data<sup>[31]</sup> |
| <u>Baculovirus release</u> |  |  |  |  |  |
| $k_{rel}$ | 9.8 | (8.3, 11.3) | $PFU \text{ cell}^{-1} h^{-1}$ | Budded baculovirus release kinetic constant | From literature<sup>[30]</sup> |
| $\tau_{rel,on}$ | 18 | (16, 20) | hpi | Budded baculovirus release onset time | From literature<sup>[30]</sup> |
| $\tau_{rel,off}$ | Cell death | – | – | End of budded baculovirus release | From literature<sup>[30]</sup> |
| <u>Transcription</u> |  |  |  |  |  |
| $k_{\Delta IE1}$ | 60.79 | (60.41, 61.17) | $nt \text{ cell}^{-1} h^{-1}$ | $\Delta IE1$ promoter transcription kinetic constant | Estimated from literature data<sup>[42]</sup> |
| $k_{polh}$ | 607.89 | (604.10, 611.68) | $nt \text{ cell}^{-1} h^{-1}$ | polh promoter transcription kinetic constant | Estimated from literature data<sup>[39]</sup> |
| $k_{p10}$ | 518.53 | (518.02, 519.04) | $nt \text{ cell}^{-1} h^{-1}$ | p10 promoter transcription kinetic constant | Estimated from literature data<sup>[39]</sup> |
| $\tau_{\Delta IE1,on}$ | 8 | (6, 10) | hpi | $\Delta IE1$ promoter transcription onset | From literature<sup>[12]</sup> |
| $\tau_{polh,on}$ | 18 | (16, 20) | hpi | polh promoter transcription onset | From literature<sup>[12]</sup> |
| $\tau_{p10,on}$ | 15 | (14, 16) | hpi | p10 promoter transcription onset | From literature<sup>[12]</sup> |
| $\tau_{transcr,off}$ | 38 | (35, 41) | hpi | Time of transcription halt for all promoters | Estimated from literature data<sup>[39]</sup> |
| <u>Translation</u> |  |  |  |  |  |
| $k_{transl}$ | $2.93 \times 10^9$ | $(2.15 \times 10^9, 3.71 \times 10^9)$ | nt cell<sup>-1</sup> h<sup>-1</sup> | Translation kinetic constant | Estimated from literature data<sup>[44]</sup> |
| $K_{transl}$ | $2.73 \times 10^4$ | $(1.20 \times 10^4, 4.26 \times 10^4)$ | mRNA cell<sup>-1</sup> | Michaelis-Menten constant for translation | Estimated from literature data<sup>[44]</sup> |
| <u>Vector genome amplification</u> |  |  |  |  |  |
| $k_{ampl}$ | $3.72 \times 10^7$ | $(1.23 \times 10^7, 6.21 \times 10^7)$ | nt cell<sup>-1</sup> h<sup>-1</sup> | Vector genome amplification kinetic constant | Estimated from literature data<sup>[44]</sup> |
| $K_{ampl}$ | $9.48 \times 10^6$ | $(6.28 \times 10^6, 1.27 \times 10^7)$ | Rep78 cell<sup>-1</sup> | Michaelis-Menten constant for vector genome amplification | Estimated from literature data<sup>[44]</sup> |
| <u>Vector genome encapsidation</u> |  |  |  |  |  |
| $k_{encaps}$ | $8.25 \times 10^2$ | $(3.80 \times 10^2, 1.27 \times 10^3)$ | nt LS<sup>-1</sup> h<sup>-1</sup> | Vector genome encapsidation kinetic constant | Estimated from literature data<sup>[44]</sup> |
| $K_{encaps,coef}$ | 5 | – | Rep52 cell<sup>-1</sup> | Coefficient for Michaelis-Menten constant for encapsidation | Fixed |
| <u>Degradation</u> |  |  |  |  |  |
| $k_{d,BV}$ | $7 \times 10^{-3}$ | $(1 \times 10^{-3}, 1.3 \times 10^{-2})$ | h<sup>-1</sup> | Extracellular baculovirus degradation kinetic constant | From literature<sup>[30]</sup> |
| $k_{d,DNA}$ | 0.01 | – | h<sup>-1</sup> | Non-encapsidated vector genome degradation kinetic constant | Estimated from literature data<sup>[47]</sup> |
| $k_{d,GFP}$ | $3.2 \times 10^{-3}$ | $(2.8 \times 10^{-3}, 3.6 \times 10^{-3})$ | h<sup>-1</sup> | GFP degradation kinetic constant | Estimated from literature data<sup>[47]</sup> |
| $k_{d,rep}$ | 0.03 | (0.02, 0.04) | h<sup>-1</sup> | Rep proteins degradation kinetic constant | Estimated from literature data<sup>[42,46]</sup> |
| $k_{d,mRNA}$ | 0.0610 | (0.0597, 0.0623) | h<sup>-1</sup> | mRNA degradation kinetic constant | Estimated from literature data<sup>[39]</sup> |

## Baculovirus Infection Dynamics

### Binding

Baculovirus binding to insect cells is a fast process at MOIs (MOI between 0.01 and 100) and cell densities (≳ 5 *×* 10^5^ cells/mL) of practical interest for the BEVS.^28,36^ The viral uptake rate of infected cells decreases as the infection cycle progresses, and eventually is completely halted due to the down-regulation of receptors induced by the virus.^28,31^ Viral binding inhibition dynamics is important for the TwoBac and ThreeBac systems, where rAAV production requires the host cell coinfection with at least, respectively, 2 and 3 different baculoviruses (one per type). Using data reported by Nielsen,^31^ we estimate the binding kinetic constant for uninfected cells (*k*_*bind*0_ = 6.3 *×* 10^*−*7^ *±* 1.1 *×* 10^*−*7^ mL cell^*−*1^h^*−*1^), and that the binding kinetic constant for infected cells (*k*_*bind*_) starts decaying exponentially at the infection time *τ*_*bind*_ = 1.8 *±* 0.5 hours post infection (hpi), with decay constant *β*_*bind*_ = 0.149 *±* 0.029 h^*−*1^. As a result, the baculovirus uptake rate from infected cells decays rapidly with the progress of infection, and infected cells do not practically bind any more virus by the time budded baculovirus is released (*τ*_*rel*_ = 18 *±* 2 hpi).^28,30,37^ Hence, budded virions do not bind to the cells that produced them, or to synchronously infected cells.

The estimated parameters are consistent with fast baculovirus binding, as previously found in the literature.^28^ According to the model, for cell concentrations larger than approximately 1 *×* 10^6^ cell mL^*−*1^, all infective virions are internalized by the cells in 4–6 hours from the process onset (not shown). As a result, for MOI as low as 3 per each baculovirus type, more than 85% and 90% of cells achieve a productive infection configuration in, respectively, ThreeBac and TwoBac systems (Fig. 2a).

**Figure 2:**
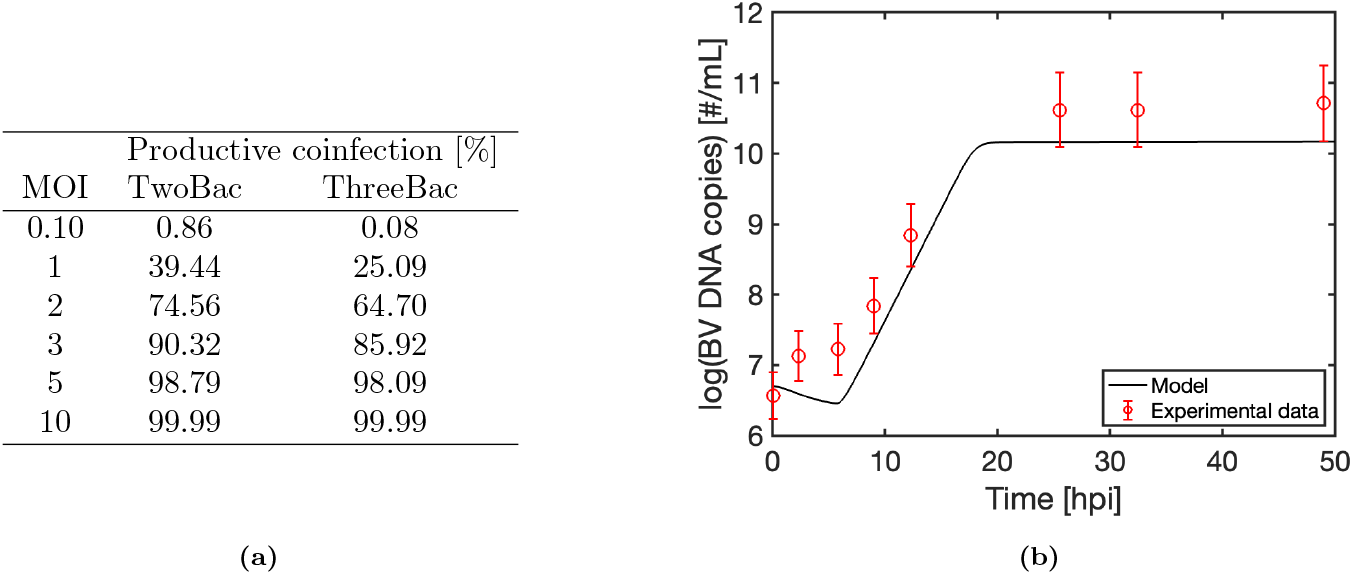
Baculovirus infection dynamics. (a) Percentage of cells that achieve productive coinfection at different MOIs. (b) Validation of cumulative extracellular and intracellular baculovirus DNA concentration: model prediction vs. experimental measurements (data from Fig. 2 of Vieira et al.).^39^

### Transport to nucleus and replication

The internalization and transport to the nucleus of receptor-bound baculovirus has been studied by Dee and Shuler,^28^ who estimated the kinetic constants for endocytosis, release into the cytosol, and transport to nucleus of baculovirus. We find that all the steps that the receptor-bound baculovirus undergoes until nuclear entry can be approximated with negligible error as a single step with kinetic constant corresponding to the rate-determining step, namely, the step presenting the lowest kinetic constant: the release of the baculovirus from the endosome into the cytosol (*k*_*int*_ = 0.6 *±* 0.12 h^*−*1^).^28^ We account for baculovirus degradation in lysosomes during trafficking to nucleus by considering that only *η* = 50% of the internalized baculovirus reaches the nucleus.^28^

The kinetic parameters for baculovirus DNA replication are estimated from data collected in four batch experiments at different cell density reported in the literature,^38^ with the model fit shown in Fig. **??**. Consistent with the literature on baculovirus infection,^12^ baculovirus DNA replication is found to start at *τ*_*repl,on*_ = 6 *±* 1 hpi, and last until *τ*_*repl,off*_ = 18 *±* 2.5 hpi. The replication kinetic constant (*k*_*repl*_) is estimated to be 0.7303 *±* 0.2382 h^*−*1^, corresponding to a baculovirus DNA doubling time of about 1 h.

### Validation of baculovirus infection dynamics

The baculovirus infection component of the model is validated by comparing the model prediction of the total intracellular and extracellular baculovirus DNA copies against the experimental measurements during a batch infection reported by Vieira and coworkers (Fig. 2b).^39^ In the first stage of the infection, baculoviruses pass from the extracellular medium to the intracellular environment, where they are, in part, degraded in lysosomes. As a result, the total intracellular baculovirus DNA level slightly decreases, until DNA replication starts at *τ*_*repl,on*_. Soon after the replication onset, the nascent baculovirus DNA becomes predominant over the baculovirus DNA that entered the nucleus following cell binding and trafficking. The baculovirus DNA level increases during replication up to five orders of magnitude, increasing from the few units that primarily infected each cell up to 10^4^–10^5^ baculovirus DNA copies per cell. The stochastic biological variability of *τ*_*repl,off*_ has a larger influence on the baculovirus DNA level at the end of replication than the baculovirus DNA level at the replication onset (Figs. 2b and **??**). The time at which replication stops practically coincides with the onset of the very late stage, during which budding occurs. Only a small fraction (1–10%) of the replicated baculovirus DNA is released as budded virus,^12,28,38^ but it is often sufficient to propagate the infection to (potentially remaining) uninfected cells in the culture medium.

## AAV Production Dynamics

### Transcription and translation

The model considers the expression through the intermediate steps of transcription and translation of the AAV structural proteins and of the non-structural proteins Rep52 and Rep78, whose genes are carried in the recombinant baculoviruses. If baculovirus promoters are included in the ITR/GOI cassette of the goiBV,^18,19^ the model can account for the expression of the transgene, as an additional protein that competes with the structural and non-structural AAV proteins for the host translation machinery.

The promoters commonly used in the BEVS for expressing the AAV proteins for rAAV production (Fig. 1a) are the *Autographa californica* polh and p10 promoters^9,18,19^ and the IE1 and ΔIE1 promoters derived from a related baculovirus, *Orgyia pseudotsugata* multiple nuclear polyhedrosis virus (OpMNPV).^18^ While the polh and p10 promoters are active in the very late infection stage, the IE1 promoter and its partially deleted form ΔIE1 present an immediate-early transcription onset, as described in the literature.^12^ The transcription kinetics for polh and p10 and the mRNA degradation kinetics are estimated by fitting the model predictions to experimental measurements of mRNA concentration in two experiments on virus-like particle production with the BEVS (Fig. **??**). Consistent with the literature,^12,19,29,40^ we find that the polh promoter (*k*_*polh*_ = 607.89 *±* 3.79 transcribed nucleotides per hour per cell, nt h^*−*1^cell^*−*1^) leads to slightly stronger transcription than the p10 promoter (*k*_*p*10_ = 518.53 *±* 0.51 nt h^*−*1^cell^*−*1^), and that the transcription onset from polh (*τ*_*polh*_ = 18 *±* 2 hpi) is delayed a few hours with respect to p10 (*τ*_*p*10_ = 15 *±* 1 hpi). Transcription from both polh and p10 promoters halts approximately simultaneously at *τ*_*transcr,off*_ = 38 *±* 3 hpi, consistent with past reports^29^. After that transcription halts, the mRNA degrades with a half-life of about 10 h (*k*_*d,mRNA*_ = 0.061 *±* 0.0013 h^*−*1^). For the ΔIE1 promoter, we set *τ*_Δ*IE*1_ = 6 *±* 2 hpi,^41^ and approximate the transcription kinetic constant as *k*_Δ*IE*1_ = *k*_*polh*_*/*10 = 60.79 nt h^*−*1^cell^*−*1^, based on an estimate from the literature^18,42^ of the relative expression rate between ΔIE1 and polh.

The model predictions are validated against mRNA concentration measurements from a dataset available in the literature^43^ in Fig. 3a. The data are from a BEVS in which a recombinant protein expression is transcriptionally regulated by the polh promoter. Baculovirus infection is carried out at low MOI, generating two infection waves. The model successfully predicts increasing target transcript concentration during the first wave. A much larger transcript concentration increase is registered in correspondence of the polh transcription phase of the second wave, namely from about 36 (= *τ*_*rel*_ + *τ*_*polh*_) to about 56 (= *τ*_*rel*_ + *τ*_*transcr,off*_) hours from the process onset (considered the time at which infectious baculoviruses are added to the system).

**Figure 3:**
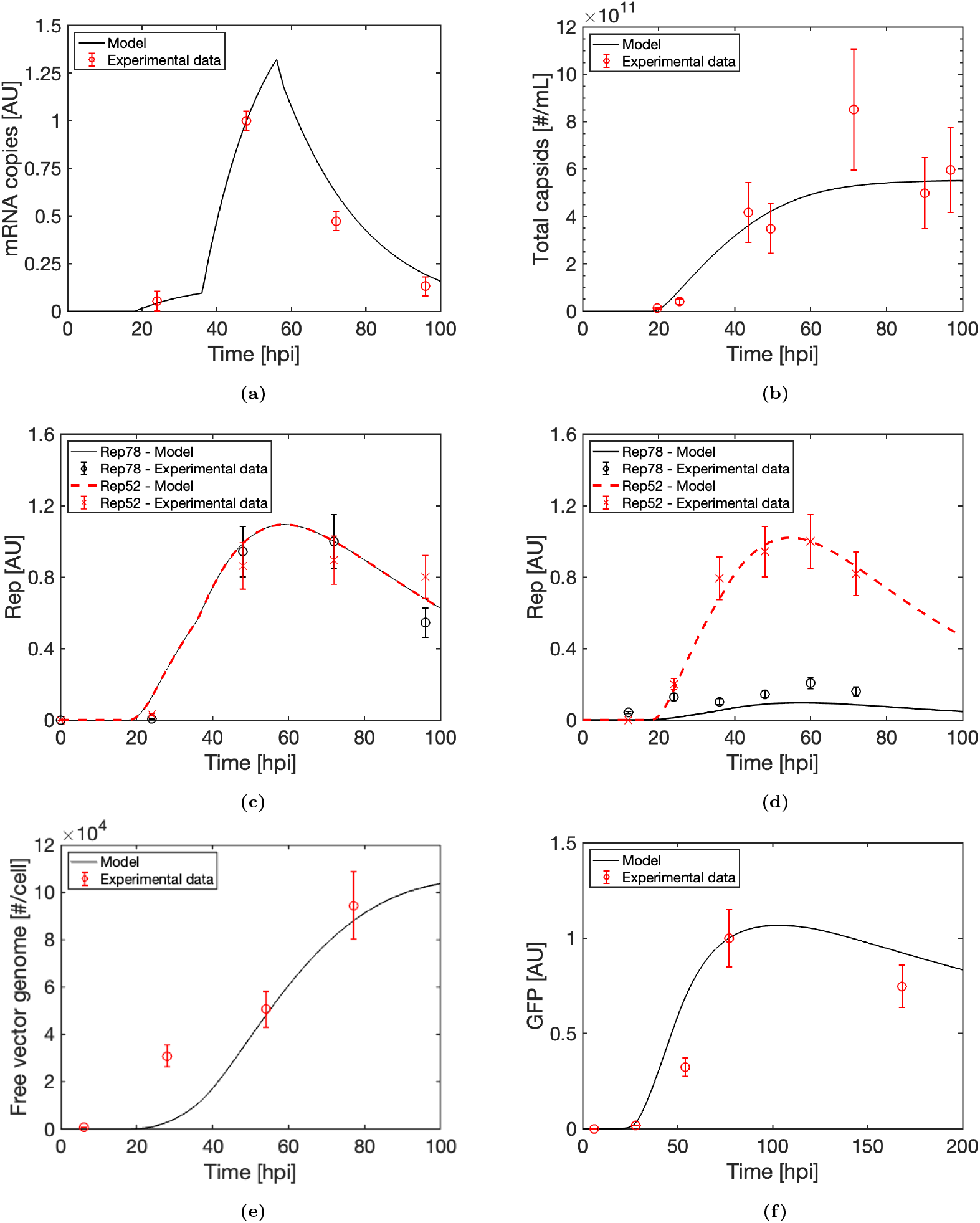
Model validation: intermediates dynamics. (a) mRNA concentration. Data from Fig. 7 (AcNPV-Bgal) of Mitchell-Logean et al.^43^ (b) Total rAAV capsids (rAAV-2). Data from Fig. 5 of Meghrous et al.^42^ (c) Rep: TwoBac. Data from Fig. 2 of Smith et al.^19^ (d) Rep: ThreeBac. Data from Fig. 4 of Urabe et al.^41^ (e) Non-encapsidated vector genome. Data from Fig. 3a of Li et al.^47^ (f) GFP. Data from Fig. 3b of Li et al.^47^

Kinetic parameters for the translation rate of the *Rep* and *Cap* transcripts are estimated (simultaneously to vector genome amplification and encapsidation kinetic parameters, see Parameter Estimation Strategy in Materials and Methods) using measurement of capsids and rAAV production from nine batch experiments with the ThreeBac system reported in the study by Aucoin et al.^44^ The employed experimental dataset is particularly convenient for this parameter estimation exercise, as the 9 batches are carried out at different MOI combinations for the three baculoviruses, providing the model with orthogonal information. The dataset contains the transducing particles concentration, as determined by gene transfer assay, rather than the filled capsids concentration. Before model calibration, we convert the infective particle concentration into filled capsid concentration by assuming a filled to infective particle ratio equal to 1000*±*500.^19,45^ The estimated parameters do not significantly change using a larger filled to infective particle ratio equal to 2000*±*1000 (not shown). The successful model validation against quantitative PCR measurements of filled capsids concentration discussed in the next subsections (specifically in subsection “rAAV production dynamics validation”) supports the conversion ratio between infective and filled particles adopted for the calibration dataset.

The estimated cumulative maximum translation rate for AAV gene transcripts is *k*_*transl*_ = 2.93 *×* 10^9^ *±* 7.8 *×* 10^8^ nt h^*−*1^cell^*−*1^, while the estimated Michalis-Menten constant for translation is *K*_*transl*_ = 2.73 *×* 10^4^ *±* 1.53 *×* 10^4^ mRNA cell^*−*1^. These parameters mean that, according to the model, in each infected cell, the sum of the translation rate of the transcripts of (if present) *Rep, Cap*, and of the transgene cannot exceed about 2.93 *×* 10^9^ nt h^*−*1^. In the absence of competition, namely in infected cells in which only one of the genes considered by the model is present, the translation rate of the relevant transcript will correspond to half of *k*_*transl*_ when the transcript concentration is equal to *K*_*transl*_.

### Capsid synthesis

The model provides a good fit of the total capsid measurements at 72 hpi in all the experiments of the calibration dataset (Fig. 4a), which are carried out at different MOI ratios for the three baculoviruses of the ThreeBac system. The weakest fit is for the batch that starts with MOI = 1 for each baculovirus, which is the only batch in which not all the cells are infected during the first wave. For this experiment, slight measurement errors in the MOIs have a large effect on the model prediction. The experimental measurements validate the model prediction that the total capsid production is larger for batches carried out at a larger ratio between the MOI of capBV and the sum of the MOIs of repBV and goiBV (Fig. 4a). The model explains this trend based on the competition between *Rep, Cap*, and transgene for expression. For instance, in the four experiments starting with MOI = 9 for capBV, the largest capsid production is achieved when MOI = 1 (MOI = 9) for both repBV and goiBV. Conversely, the lowest capsid production is achieved when MOI = 9 for both repBV and goiBV. The capsid production of the two batches in which the sum of the MOIs of repBV and goiBV is 10 (i.e., MOI = 9 for repBV and MOI = 1 for goiBV, and vice versa) lies between these two extremes.

**Figure 4:**
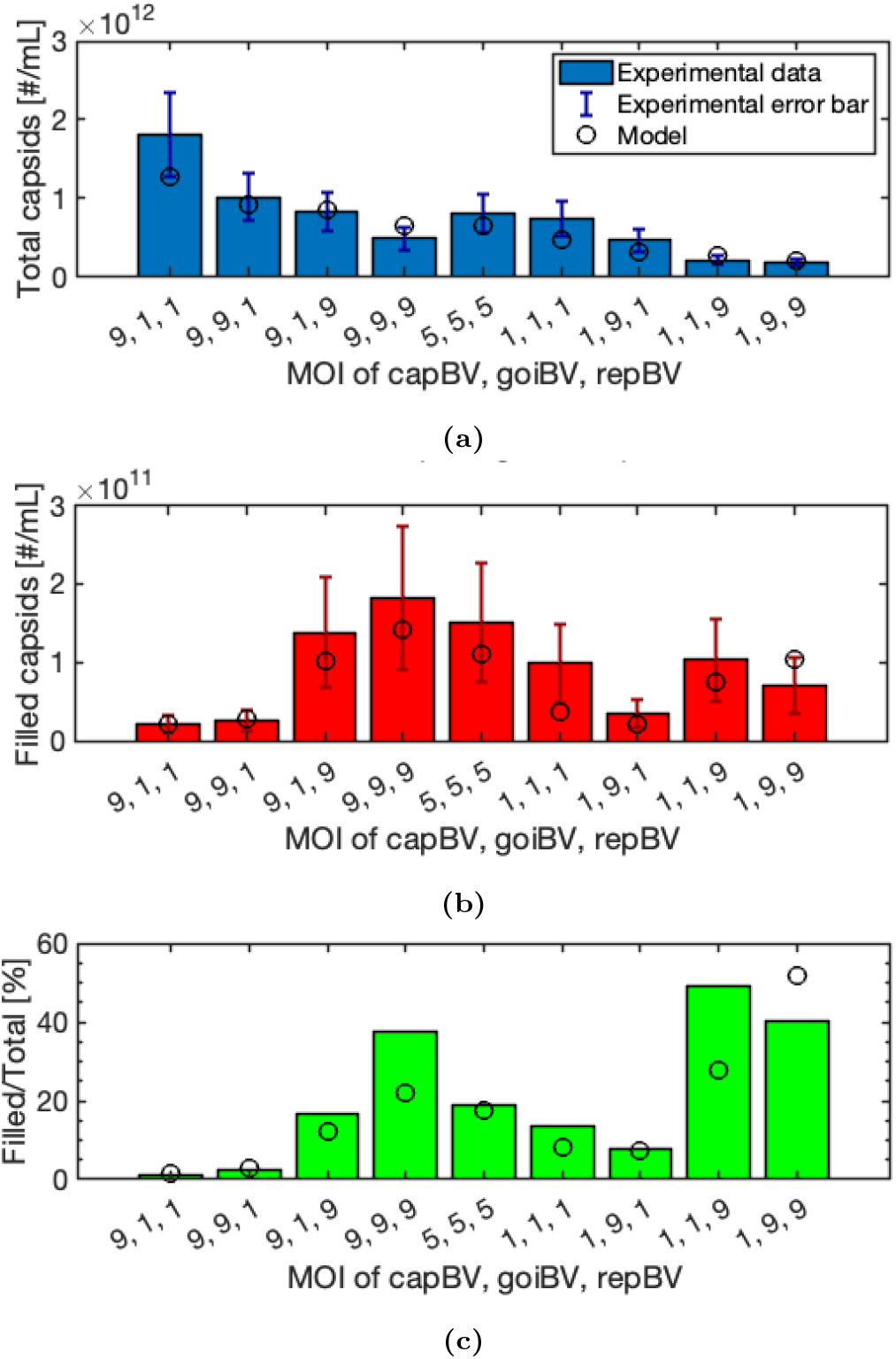
Model fit to the ThreeBac dataset (rAAV-2) used for estimation of *k*_*transl*_, *K*_*transl*_, *k*_*ampl*_, *K*_*ampl*_, and *k*_*encaps*_: model predictions (circles) and experimental data (bars). Error bars refer to experimental data. (a) Total rAAV capsids at 72 hpi. Data from Fig. 3 of Aucoin et al.^44^ (b) Filled rAAV capsids at 96 hpi. The reported experimental data are obtained by multiplying the infective viral particle titer reported in Fig. 2a of Aucoin et al. by a factor of 1000. (c) Filled to empty capsids ratio.

We additionally validate the model prediction of the capsid concentration against experimental measurements from another dataset reported in the literature for rAAV production in the ThreeBac system (Fig. 3b). The combined MOI (= 5) used in the experiment is sufficiently large to infect all cells with one wave. The capsid concentration profile predicted by the model is well aligned to experimental measurements. No capsids are present in the system until about 20 hpi, shortly after that the polh promoter of the *Cap* cassette becomes active (*τ*_*polh*_ = 18 h). The capsid concentration rises quickly until about 40 hpi. At that point, the capsid synthesis rate decreases due to a halt in transcription (*τ*_*transcr,off*_ = 38 h) and a loss of cell viability. Almost no more capsids are produced after 72 hpi, due to low mRNA levels (Fig. 3a) and viability. No significant capsid degradation is registered, validating the model hypothesis.

### Rep protein synthesis

The model predictions of Rep52 and Rep78 levels in TwoBac and ThreeBac systems are validated by experimental datasets from the literature (Figs. 3cd). The good agreement between model predictions and measurements is particularly interesting for TwoBac, since the model parameters for *Rep* expression have been estimated on data collected in the ThreeBac system.^44^ This result validates the model hypothesis that the dynamics of transcription from the *Rep52* polh promoter in ThreeBac and from the *Rep* polh promoter in TwoBac are the same, and that the translation dynamics is equivalent in the two systems.

In the TwoBac process (Fig. 3c), Rep52 and Rep78 are present at similar levels.^19^ The level of Rep proteins starts rising around 20 hours from the process onset, shortly after the initiation of transcription from the polh promoter of the *Rep* cassette, and peaks at about 60 hours from the process onset, corresponding with the end of transcription for the second wave of infected cells. After that point, Rep proteins degrade with a half-life of about 20 h (we estimate *k*_*d,rep*_ = 0.03 h^*−*1^ using data from the literature^42,46^).

In the ThreeBac process, the Rep52 level follows the same trend as in TwoBac (Fig. 3d), since *Rep52* is expressed through the polh promoter also in this case (Fig. 1). The level of Rep78 is much lower, as its expression is regulated by the weaker Δ*IE*1 promoter. The earlier onset of transcription from Δ*IE*1 with respect to polh is not enough to compensate for the difference in strength of the promoters. Actually, low baculovirus DNA level is registered before 18 hpi (Fig. 2b), when transcription is active from Δ*IE*1, but is not active from polh.

### Vector genome amplification

Rep78 activities are critical for vector genome amplification.^7^ Hence, vector genome amplification cannot start until Rep78 has been expressed, and the overall process can be limited by a Rep78 deficiency.^26^ To directly estimate the extent of this limitation in the BEVS, we implement vector genome amplification in our model through a Michaelis-Menten kinetics. We estimate, from the calibration dataset,^44^ that the maximum vector genome amplification rate (in the absence of Rep78 limitation) is *k*_*ampl*_ = 3.72 *×* 10^7^ *±* 2.49 *×* 10^7^ nt cell^*−*1^ h^*−*1^. The Michaelis-Menten constant is equal to *K*_*ampl*_ = 9.48 *×* 10^6^ *±* 3.20 *×* 10^6^ Rep78 cell^*−*1^ (number of Rep78 proteins per cell). Experimental data^47^ show that non-encapsidated vector genome degrades with a half-life of about 70 h (*k*_*d,DNA*_ = 0.01 h^*−*1^).

We validate the vector genome amplification kinetics predicted by the model against experimental measurements from the literature.^47^ Data was collected during a repBV infection of insect cells stably transfected with an ITR-flanked GFP (Fig. 3e). No vector genome was encapsidated during the experiment, since *Cap* was not introduced in the host. Even though the model parameters were estimated from ThreeBac experiments, the model accurately predicts the experimentally measured maximum level of vector genome in the stable cell line infection (*≈* 8 *×* 10^4^ copies per cell, achieved about 3 days post infection, dpi). The (non-encapsidated) vector genome copy number starts growing at about 25 hpi, corresponding to the time at which the level of Rep78 starts to increase during a repBV infection (Fig. 3d). The vector genome copy number peaks at about 80 hpi and then starts declining, due to DNA degradation. The ITR-flanked GFP cassette stably transfected in the cell line used in the experiment contains a p10 promoter, for transgene expression in insect cells. The measured concentration of GFP during the experiment is well predicted by the model (Fig. 3f). As soon as the vector genome is amplified in sufficient numbers (i.e., at about 25 hpi), GFP is detected in the system. Mimicking vector genome amplification dynamics, the GFP synthesis rate peaks about 3 dpi.

Additional validation of the vector genome amplification rate predicted by the model is in Fig. **??** for an additional literature dataset.^41^

### Vector genome encapsidation

Vector genome encapsidation is the last step of rAAV particle formation. From the calibration dataset,^44^ the encapsidation kinetic constant is estimated to be *k*_*encaps*_ = 8.25 *×* 10^2^ *±* 4.45 *×* 10^2^ nt h^*−*1^ LS^*−*1^, where LS (= limiting species for encapsidation) is the species between empty capsids and non-encapsidated vector genome that has lower concentration in the cell (Eq. 36).

The model provides a good fit of the filled capsid production at the end of the batch for the nine ThreeBac experiments from the literature used for model calibration (Fig. 4b). The model simulation also resembles the intermediate data collected during the experiments, not used for model calibration (Fig. **??**). The largest rAAV production is achieved in the experiment carried out at the highest MOI (= 9) for all the three baculoviruses, according to both the model and the experimental measurements. All the experiments for which a MOI = 9 was used for repBV have similar rAAV production, independently from the MOIs of capBV and goiBV, highlighting the important role of Rep proteins for the production of filled capsids. The model predicts a small increase of full rAAV for experiments that have a larger MOI of goiBV (in the same MOI conditions for capBV and repBV).

The percentage of filled particles (Fig. 4c) increases significantly with the ratio between the MOI of repBV and the MOI of capBV. For instance, the two batches with lowest ratio between repBV MOI and capBV MOI (= 1*/*9), have the lowest ratio of filled capsids, according to both model and experimental measurements, even though they lead to the largest total capsid production. The two batches with the largest ratio between repBV MOI and capBV MOI (= 2) have the largest ratio of filled to total capsids. The percentage of filled particles also modestly increases with the MOI of goiBV (at fixed MOI for capBV and repBV).

### Validation of rAAV production dynamics

Model validation for the baculovirus infection compartment (Fig. 2b) and for the intermediates of rAAV production (including mRNA, capsids, Rep proteins, and non-encapsidated vector genome; Fig. 3) is discussed above. This section describes the successful validation of the overall baculovirus infection and rAAV production model, by comparing the model prediction of the filled capsid concentrations to measurements from additional ThreeBac, TwoBac,^11,19^ and OneBac^9^ literature datasets. In all the considered datasets, the filled capsids concentration is accurately quantified through PCR.

First, consider a study carried out in the ThreeBac system.^46^ The model simulation is well-aligned with the experimental measurements (Fig. 5a). A low total MOI (= 0.3), compatible with a two-wave infection, was used in the experiment. Any rAAV produced from the first wave of infected cells was below the limits of detection. The filled capsid concentration starts to grow about 45–50 h from the process onset, namely 25–30 hpi for cells infected in the second wave, with a delay of about 10 h from the onset of capsid synthesis (Fig. 3b). Then the filled capsid concentration increases steadily, until it stabilizes about 4.5 days from the process onset, due to loss of viability.

**Figure 5:**
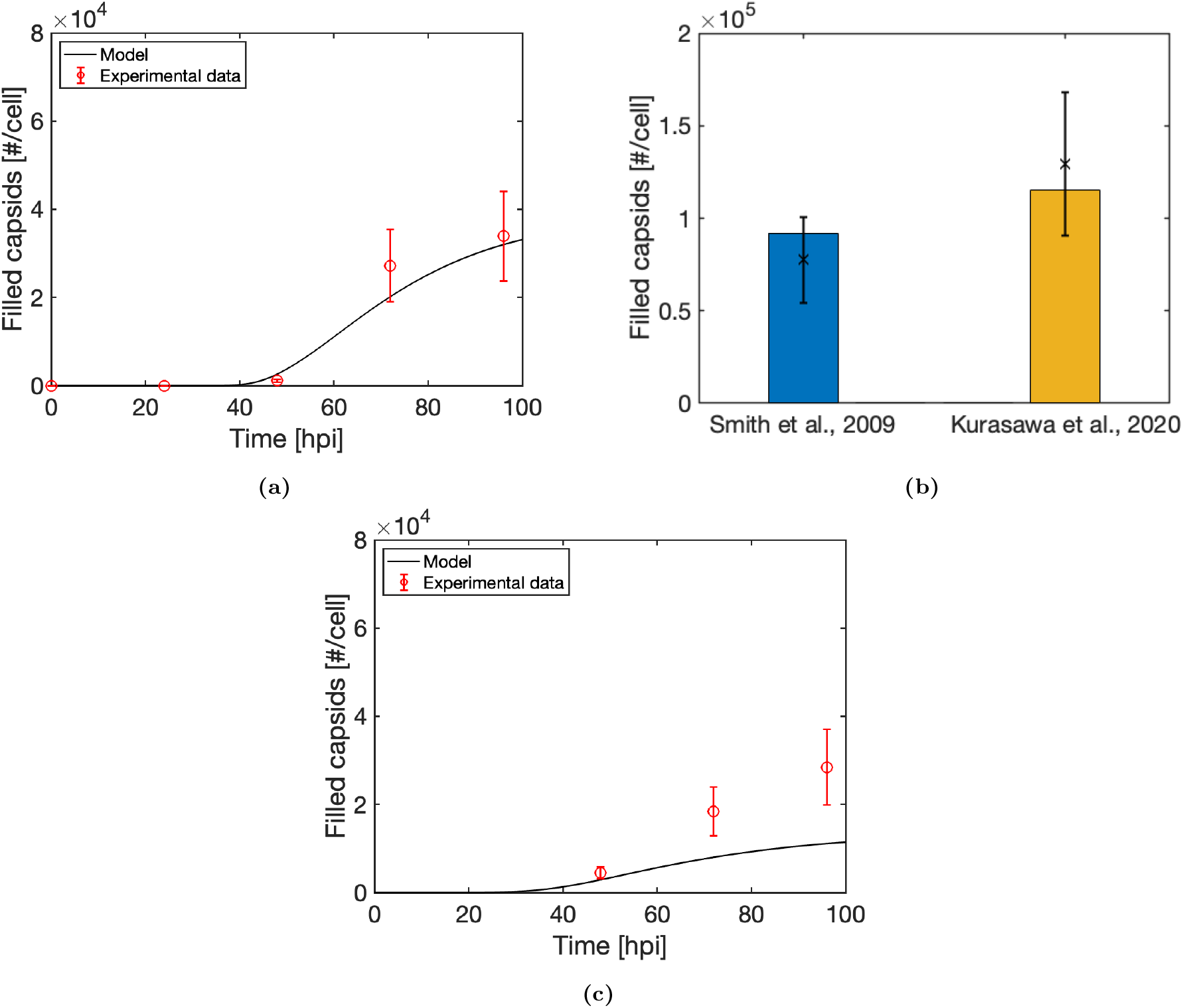
Model validation: rAAV production. (a) ThreeBac. Data (rAAV-2) from Fig. 5 of Mena et al.^46^ (b) TwoBac. Data (rAAV-2) from Table 1 of Smith et al.^19^ and from Figure 3a of Kurasawa et al.^11^. Model predictions (bars) and experimental data (markers). (c) OneBac. Data (rAAV-5) from Fig. 4a (bioreactor) of Joshi et al.^9^

All the data discussed so far with respect to model calibration and validation refer to experiments in the ThreeBac system (except for the validation of Rep proteins profiles in TwoBac, Fig. 3c). Without any change to the estimated parameters, the TwoBac implementation of the model predicts, within the measurement error, the filled capsids production in two studies that used the TwoBac system^11,19^ (Fig. 5b). As predicted by the model, the filled capsid production per cell achieved by Smith et al.^19^ is lower than obtained by Kurasawa et al. ^11^ despite both experiments being carried out in similar conditions, including initial MOIs. The main difference between the two experimental procedures lies in the goiBV construct. While Smith et al. used the goiBV reported in Fig. 1a, the goiBV used by Kurasawa et al. had a 5% shorter ITR-flanked sequence, and included only the CMV promoter (no p10 promoter). Different viability profiles (not fully reported by the authors) or lab-to-lab variability might play a role in the production difference between the two studies. However, the model fully explains the increase in rAAV production based on the difference between the goiBVs used in the two studies. According to the model, the absence of the strong p10 promoter in the ITR-flanked sequence of goiBV leads, by itself, to about a 20% increase of filled capsid production, due to an increase of *Rep* and *Cap* expression as a consequence of reduced competition with transgene expression (as discussed for capsids production; Fig. 4a). In addition, the model also estimates that the shorter ITR-flanked sequence can be amplified faster, leading to a corresponding production increase of 5%.

Differently from TwoBac and ThreeBac, the *Rep* and *Cap* cassettes in the OneBac system^9^ are stably integrated in the genome of the insect cells, and rAAV is produced upon infection using only one baculovirus that carries the ITR-flanked transgene.^20^ Recently, Joshi et al.^9^ achieved rAAV-5 titers (2.5 *×* 10^11^ *−* 3.5 *×* 10^11^ filled capsids per mL) with OneBac higher than previously demonstrated with TwoBac in 1L-scale bioreactors for several rAAV serotypes.^10,48^ Figure 5c compares the prediction of the OneBac implementation of the model against measurements of filled capsid concentration for a representative experiment from the study of Joshi et al.^9^ Although the total rAAV production is underestimated, the model successfully describes the main rAAV production dynamics (Fig. 5c), and predicts that about 20% of the produced capsids are filled, as found experimentally.^9^ This validation, carried out without any re-estimation of the model parameters, is particularly challenging, because the *Rep* and *Cap* integrated in the cells of OneBac have many differences with respect to those in the experiments used for model calibration.^49^ For instance, the Kozak consensus sequences are different, and *Cap* comes from AAV-5, rather than AAV-2 as in the experiments considered for model calibration. Interestingly, the rAAV production per cell of the considered OneBac experiment is comparable to that of ThreeBac (Fig. 5a), and is 3–4 times lower than the production per cell achieved with TwoBac (Fig. 5b). The high cell density (*≈* 1 *×* 10^7^ cell/mL) to which cells have been grown in the study by Joshi et al.^9^ is the main factor that led to the high titer obtained with OneBac, rather than a greater productivity per cell compared to TwoBac.

### Cell Growth and Death Dynamics

Uninfected Sf9 cells have a growth kinetic constant (*μ*) of 2.8 *×* 10^*−*2^ h^*−*1^,^30,46^ and a low death kinetic constant (*k*_*death,T*_) of about 8 *×* 10^*−*5^ h^*−*1^.^30^ In turn, baculovirus-infected Sf9 cells do not undergo mitosis, and have an increased death rate that leads to death of the whole population of synchronously infected cells in 3–5 dpi.^12^

It has been suggested that the death rate of infected cells is correlated to the concentration in the host of baculovirus DNA,^29^ of Rep78 (known to cause cytotoxicity in mammalian cells^50^), and of rAAV capsids (due to potential cytolytic effect). We use viability measurements (Fig. **??**) across the nine batch experiments with the ThreeBac system carried out by Aucoin et al.^44^ to estimate the correlation of the death rate of infected cells with the model estimations of the levels of baculovirus DNA, Rep78, and total rAAV capsids in the cell. The infected cell viability shows no significant decay until a certain time after infection (*τ*_*death*_), corresponding to 24 *±* 2 hpi. After that, the death rate is correlated to the logarithm of the baculovirus DNA concentration, proportionally to *k*_*death,DNA*_ = 2.9 *×* 10^*−*3^ *±* 4 *×* 10^*−*4^ h^*−*1^. Interestingly, the parameter estimation indicates that the death rate is not directly influenced by the concentration of Rep78 (*k*_*death,rep*_ = 0 h^*−*1^). The dependency of the death rate on the rAAV capsid concentration is also found to be negligible (*k*_*death,cap*_ = 0)h^*−*1^), as it does not improve the model fit to the cells viability in a statistically significant way, according to the Akaike information criterion.^51^ The estimated parameters are generally representative of the death rate dynamics for baculovirus-infected insect cells.^12,30^ These results indicate that the cell death rate is directly correlated to the baculovirus DNA concentration, while there is no direct correlation between cell death rate and the concentration of Rep78 and capsids. Hence, the accelerated death rate experienced by infected cells is mainly due to the expression of baculovirus proteins, and rAAV production does not diminish cell viability.

### In-silico Analysis of the Process

We use the validated model for baculovirus infection and rAAV production to gain insights into the process dynamics. The focus is on TwoBac and ThreeBac, for which we carried out a comprehensive model validation, also with respect to the intermediates. Sample simulation plots for MOI = 3 for each baculovirus and cell density of 2 *×* 10^6^ cells ml^*−*1^ for TwoBac and ThreeBac are in Fig. 6. The corresponding rates for the intracellular reactions that lead to rAAV formation are reported in Fig. 7a. A sensitivity analysis of the model parameters, expressing their relative impact on rAAV production for TwoBac, is shown in Fig. 7b.

**Figure 6:**
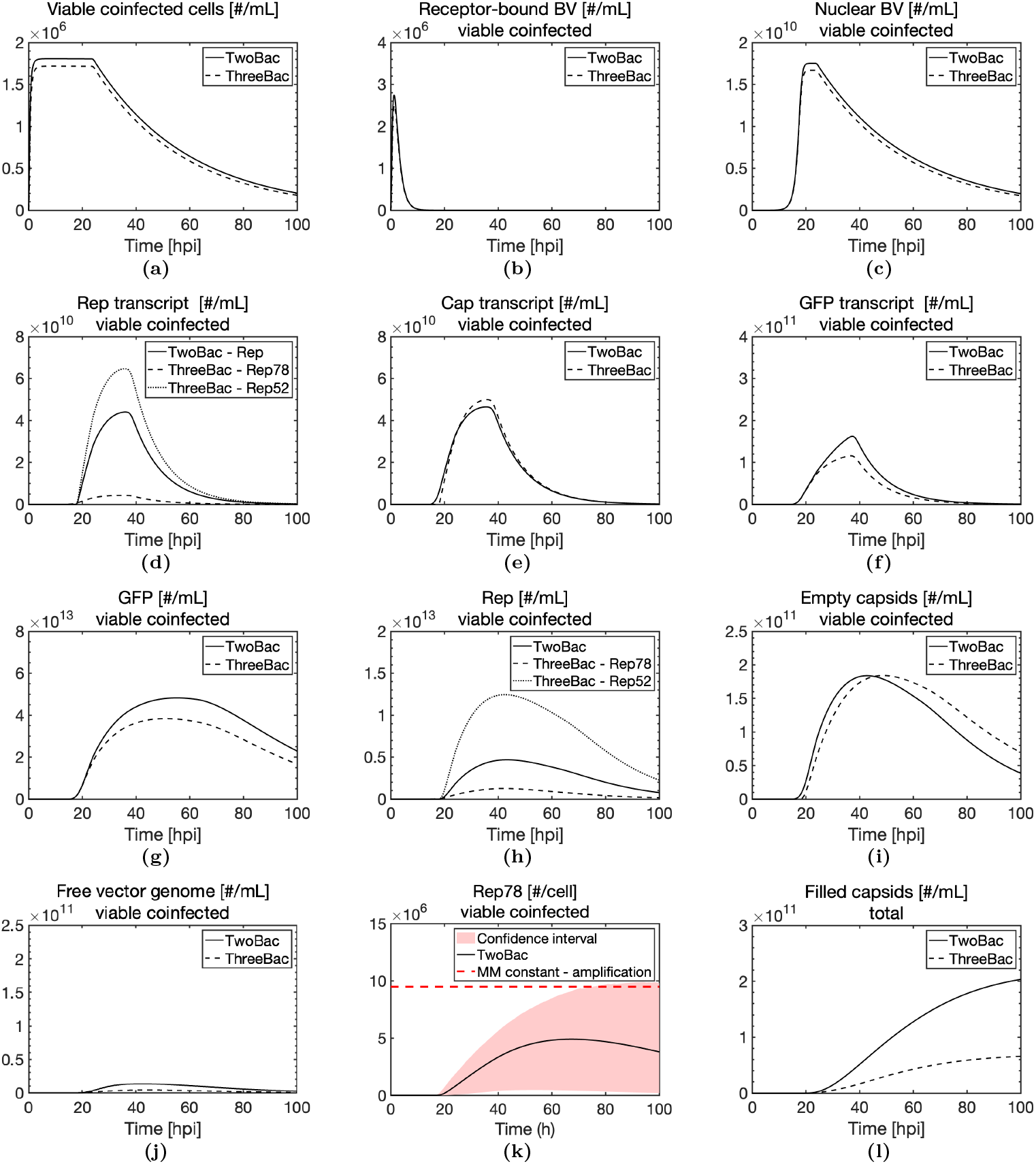
In-silico analysis of the process: sample plots for TwoBac and ThreeBac, generated for MOI = 3 per each baculovirus and cell density at infection equal to 2 *×* 10^6^ cell mL^*−*1^. (a) Viable coinfected cell density. (b) Baculovirus bound to viable coinfected cells (for each type of BV of the system). (c) Baculovirus DNA in nucleus of viable coinfected cells (for each type of BV of the system). (d) *Rep* transcripts in viable coinfected cells. (e) *Cap* transcripts in viable coinfected cells. (f) Transgene transcript in viable coinfected cells. (g) GFP in viable coinfected cells. (h) Rep proteins in viable coinfected cells. (i) Empty capsids in viable coinfected cells. (j) Free vector genome available for encapsidation in viable coinfected cells. (k) Concentration of Rep78 in coinfected cells for TwoBac: estimation and confidence interval from Monte Carlo simulation. The estimated Michaelis-Menten constant for Rep78 limitation to vector genome limitation is also reported. (l) Total concentration of filled rAAV capsids in the system. For (b–j), species concentration is reported as cumulative concentration in the batch, obtained by multiplying the level of the species in viable coinfected cells by the (local) density of viable coinfected cells.

**Figure 7:**
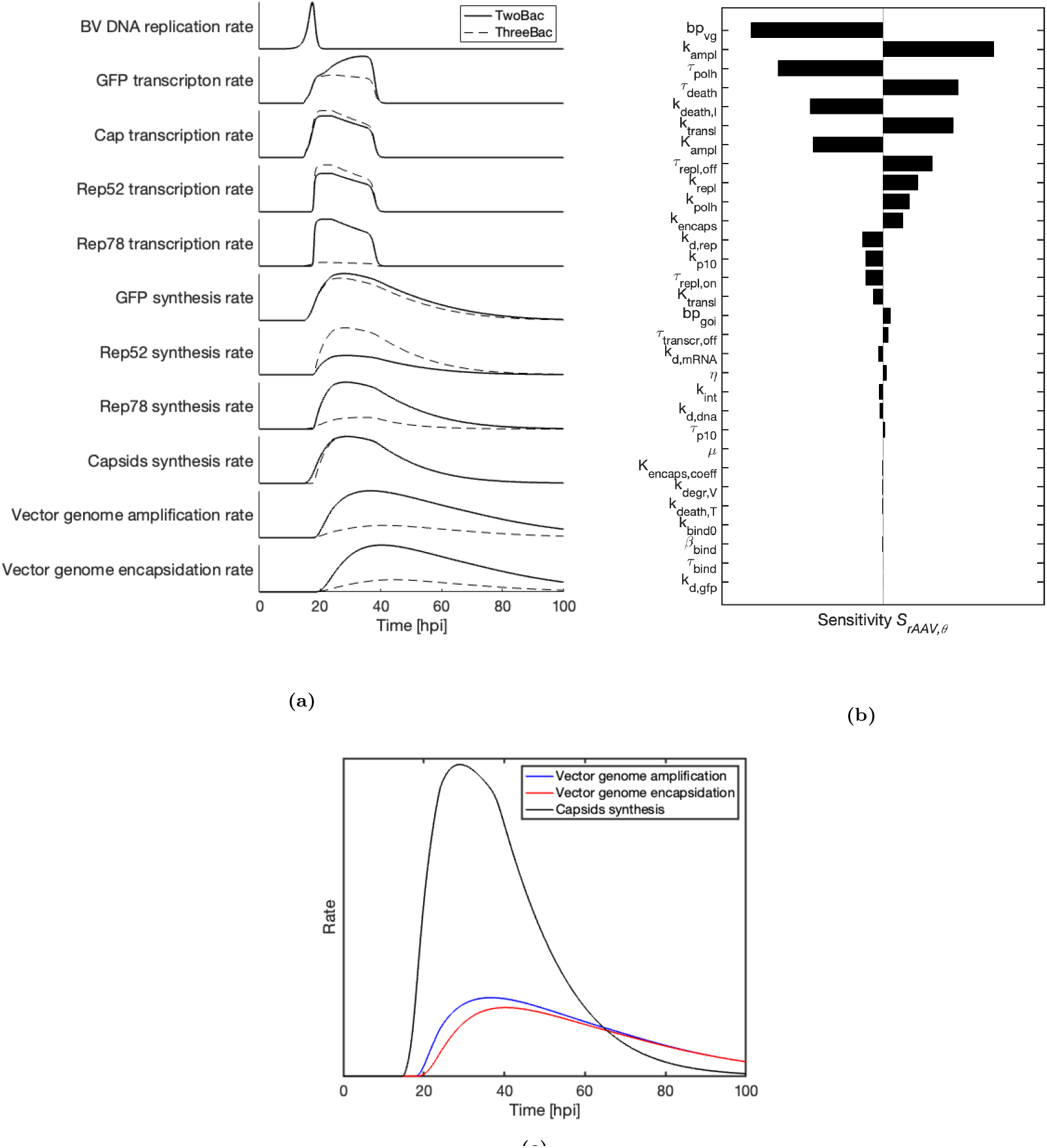
In-silico analysis of the process: reaction rates and sensitivity analysis, generated with MOI = 3 per BV and cell density at infection equal to 2 *×* 10^6^ cell mL^*−*1^. (a) Dynamic trends of reaction rates for TwoBac (continuous line) and ThreeBac (dashed line). (b) Sensitivity of rAAV production to model parameters (Eq. 49). Sensitivities with respect to *k*_*d,rep*_ and *k*_*d,rep*_ are not reported, since they are found not to affect the model response. (c) Comparison among rates of vector genome amplification, vector genome encapsidation and rAAV (empty) capsids synthesis in TwoBac simulation.

Most (*>*80%) cells achieve a productive coinfection for both TwoBac and ThreeBac (Fig. 6a). A significant viability decrease is registered starting from 24 hpi, and less than 25% are still viable 3 dpi. Figure 6b–j show the cumulative concentration in the system of the intermediates for rAAV production in viable cells having a productive coinfection. This representation allows assessments of the total concentration of the intermediates that can actually take part in rAAV formation.

The concentration of baculovirus bound to host receptors (Fig. 6b) peaks at the process onset, caused by fast baculovirus binding, followed by internalization and transport to nucleus. The copy number of baculovirus DNA in the nucleus grows moderately in the first phase of the process, due to the internalization of the extracellular baculovirus, and exponentially in the replication phase (6–18 hpi; Figs. 6c and 7a). From 24 hpi, the cumulative baculovirus DNA copy number in coinfected viable cells decreases, due to loss of cell viability, although the copy number in cells that are still viable remains approximately constant.

Transcript concentration profiles (Figs. 6d–f) are closely related to the transcription dynamics of the associated promoters. The concentration of all transcripts starts decaying significantly afterthe halt of transcription (*≈* 40 hpi), due to mRNA degradation and loss of cell viability. *Cap* transcript concentration dynamics in TwoBac and ThreeBac are similar (Fig. 6d), with slightly earlier transcription onset in TwoBac (p10 promoter) than in ThreeBac (polh promoter). A single *Rep* transcript is produced in TwoBac, while *Rep52* and *Rep78* are transcribed separately in ThreeBac (Fig. 1). Transcription of *Rep78* (ΔIE1 promoter) in ThreeBac is much weaker than *Rep* transcription (polh promoter) in TwoBac (Figs. 6d and 7a). The GFP transcript reaches a higher level in TwoBac than in ThreeBac, due to the larger amount of template available in TwoBac as a consequence of stronger vector genome amplification.

Protein levels (Figs. 6ghi) start to rise as soon as transcription of the corresponding genes is initiated (Fig. 7a). Rep52 is less abundant in TwoBac than in ThreeBac (6h), due to the lower transcript concentration and to the weaker translation caused by the leaky-scanning mechanism. On the other hand, the level of Rep78 is much higher in TwoBac than in ThreeBac. The capsid synthesis rate and the concentration of empty capsids across the batch are similar for the two systems (Figs. 6i and 7a).

The level of free vector genome available for encapsidation is much lower than the empty capsids level for both TwoBac and ThreeBac (Figs. 6ij). By overlapping the rates of capsid synthesis, vector genome amplification, and encapsidation in TwoBac (Figs. 7c), the model indicates that the encapsidation rate is controlled by the amplification rate, and that both are significantly lower than the capsid synthesis rate. Hence, non-encapsidated vector genome is a limiting species for rAAV formation, and vector genome amplification is the limiting step within the reaction-transport network. Comparing the Rep78 level in coinfected cells for the TwoBac system against the estimated Michalis-Menten constant for amplification, the model indicates that Rep78 is limiting for vector genome amplification, also when the uncertainty of the model prediction is considered (Fig. 6k). In TwoBac, encapsidation appears to be limited only by the availability of non-encapsidated vector genome. Model simulation shows that about 90% of the synthesized vector genome copies are successfully packaged, with most of the remaining genomes not being encapsidated due to loss of cell viability (Fig. **??**). For ThreeBac, the non-encapsidated vector availability is even more limiting for rAAV production (Fig. 6j), due to the lower Rep78 level (Fig. 6h) and the consequent lower vector genome amplification and encapsidation rates (Figs. 7c), compared to TwoBac.

The sensitivity analysis for TwoBac also indicates that vector genome amplification is the limiting step for rAAV production (Fig. 7b). Of all the model parameters, the maximum vector genome amplification rate (*k*_*ampl*_) affects rAAV production the most, whereas the Michaelis-Menten constant for amplification (*K*_*ampl*_) is the sixth parameter in the ranking. The time of transcription onset from the polh promoter (*τ*_*polh*_), controlling *Rep* transcription, is also significantly affecting production, together with the parameters related to viability (*τ*_*death*_ and *k*_*death,I*_). The maximum translation rate (*k*_*transl*_) is the fifth parameter in the ranking, while the rest of the parameters that have a non-negligible effect on production are related, in order of decreasing importance, to baculovirus DNA replication (*τ*_*repl,off*_, *k*_*repl*_, *τ*_*repl,on*_), transcription (*k*_*polh*_ and *k*_*p*10_), encapsidation (*k*_*encaps*_), and Rep protein degradation (*k*_*d,rep*_).

## Discussion

Achieving high productivity in rAAV manufacturing is of utmost importance for the successful delivery of gene therapy treatments to patients. Still, current rAAV manufacturing processes with both mammalian and insect cells present severe limitations in terms of productivity. In this work, we developed the first model for rAAV production in the BEVS, to shed light on the infection and intracellular dynamics that limit rAAV production in state-of-the-art processes.

### A Validated Quantitative Model for Baculovirus Infection and rAAV Production

The model predictions are in excellent accordance with experimental measurements of rAAV and of intermediates reported in the most significant studies from the literature of rAAV production with the BEVS. We demonstrated successful model validation for both TwoBac (Figs. 3c and 5b) and ThreeBac experiments (Figs. 3b, 3d, 4, 5a, **??** and **??**) and, without any additional parameter estimation, for experiments using stable cell lines with integrated ITR/GOI (Figs. 3e and 3f) or *Rep* and *Cap* (Fig. 5c) cassettes.

Regarding the baculovirus compartment, the model shows good agreement with experimentally measured baculovirus DNA concentration during infection (Figs. 2b and **??**). According to the model, productive coinfection is achieved in the vast majority of cells for MOI per baculovirus as low as 3 in both TwoBac and ThreeBac (Figs. 2a and 6a). Experimental measurements and model analysis show that, in the replication phase (*≈* 6–18 hpi), the baculovirus DNA level grows from the few copies that infected the host up to *≈* 10^5^ copies per cell (Figs. 6c and 2b). The final DNA level at the end of replication seems to be more dependent on the biological variability of *τ*_*repl,off*_ (Table 1), rather than on infection conditions such as MOI or cell density (Fig. **??**). Hence, both experimental and modeling results indicate that rAAV production is not significantly impacted by unsuccessful or unproductive coinfection or low baculovirus level, for MOI per each baculovirus as low as 2–3.

The rAAV production compartment of the model accounts for the expression of AAV genes through the intermediate steps of transcription and translation, for the synthesis of rAAV capsids and of Rep52 and Rep78 proteins, and, finally, for vector genome amplification and encapsidation. The model is conceived with a plug-and-play structure, which can simulate gene expression through different gene-promoter arrangements, such as used in TwoBac and ThreeBac (Fig. 1a). The model simulation is successfully validated against mRNA concentration measurements collected during BEVS experiments (Figs. 3a and **??**). The model predictions are also aligned with respect to experimental concentrations of proteins related to rAAV production, including the total rAAV capsid concentration (Figs. 3b and 4a), Rep proteins (Figs. 3c and 3d), and transgene protein (Fig. 3f). Model simulation and experimental results highlight competition between *Cap, Rep*, and transgene for the host translation machinery. Accordingly, the model estimates that stronger transgene expression results in reduced rAAV production, as previously found experimentally^44^ (Figs. 4 and 5b). Experimental measurements also validate the model prediction for the vector genome amplification (Figs. 3e and **??**) and encapsidation dynamics, including the predicted rAAV titer achieved in OneBac, TwoBac, and ThreeBac (Fig. 5).

### Limiting Steps for rAAV Production

Model simulation for TwoBac shows that, in coinfected cells, the concentration of free vector genome available for encapsidation (Fig. 6j) is generally much lower than the concentration of empty capsids (Fig. 6i). Hence, non-encapsidated vector genome is the limiting species for rAAV formation, up to the point that the encapsidation rate is essentially controlled by the amplification rate (Fig. 7c). The model indicates that vector genome amplification is, in turn, limited by Rep78 (Figs. 6k and 7b). According to the model, the genome limitation is stronger in ThreeBac than in TwoBac (Figs. 6ij and 7a), due to weaker *Rep78* expression (Figs. 6dh) and to consequent weaker vector genome amplification (Fig. 7a).

Experimentally, rAAV production in ThreeBac (Fig. 5a) is typically lower than in TwoBac (Fig. 5b). The literature attributes the greater rAAV production in TwoBac to a larger coinfected cell population, producing rAAV for a longer period of time prior to cell death.^22^ Instead, the model-based analysis indicates that the use in TwoBac of the polh promoter for expressing *Rep78*, rather than the weaker ΔIE1 as in ThreeBac (Fig. 1a), plays a main role in the achievement of a larger rAAV productivity. Indeed, *Rep78* expression has experimentally been found to be stronger in TwoBac (Fig. 3c) than in ThreeBac (Fig. 3d). Additional findings from the literature confirm that stronger *Rep78* expression is correlated to larger vector genome amplification and filled capsid production.^47^ Experiments on a stable cell line with integrated ITR/GOI cassette^47^ show that, upon infection with repBV, the non-encapsidated vector genome copy number reaches a significantly higher level with increasing MOI (experiments carried out at MOI = 1, 2, 3), and consequent higher level of *Rep78* templates in the cell. Additionally, the data reported by Aucoin et al.^44^ confirm that the filled-to-empty particles ratio grows with the ratio between *Rep* and *Cap* templates in the host (Fig. 4c).

The model-based finding according to which transgene expression limits rAAV production due to competition for the host translation machinery is also supported by experimental evidence (Fig. 4a). Hence, reduction of transgene expression by removing insect cell promoters from the ITR/GOI cassette in the BEVS can lead to significant increase in filled rAAV capsids (although a proportional increase of empty capsids is also expected). For rAAV manufacturing in mammalian cells, instead, it is challenging to reduce transgene expression without also losing the therapeutic effect. Following the same rationale, productivity in the BEVS can also be enhanced by reducing or suppressing expression of baculovirus genes not useful for production of rAAV or budded baculovirus.

The model indicates that encapsidation and Rep52 deficiency are not limiting rAAV production in TwoBac and ThreeBac. Model simulation for TwoBac, in which amplification is more effective (Fig. 7a) and *Rep52* is more strongly expressed (Fig. 6h), shows that roughly 90% of the vector genomes are effectively encapsidated, with the remaining 10% of vector genomes not being packaged for loss of viability (Fig. **??**). The Michaelis-Menten coefficient expressing Rep52 limitation to packaging (*K*_*encaps,coeff*_) is also found to have a negligible sensitivity on rAAV production (Fig. 7b). Experimentally, repBV infection in a stable cell line with integrated ITR/GOI leads to vector genome amplification (Fig. 3e) up to levels similar to filled capsid production in ThreeBac (Fig. 5a), once the competition between *Rep* and *Cap* for expression (occurring in ThreeBac but not in the stable cell line experiment) is accounted for. This experimental finding further supports the model-based result that encapsidation and Rep52 are not limiting filled capsid production in TwoBac and ThreeBac.

Regarding OneBac, the model suggests that the low filled capsid production per cell (Fig. 5c) is due to low vector genome amplification also in this case. Rep78 limitation to vector genome amplification emerges also for OneBac. The ratio between the integrated *Cap* and *Rep* gene copies per cell in the experiment has been measured to be about 10:1,^9^ which the model indicates is not optimal for achieving a large filled-to-empty capsid ratio (Fig. 4). However, model validation has not been carried out for OneBac as exhaustively as for TwoBac and ThreeBac, and a more detailed analysis would be needed to draw reliable conclusions on production bottlenecks in OneBac.

Maintaining high cell viability for as long as possible during baculovirus infection is crucial for achieving large rAAV titers (Fig. 7b). Model fit to viability measurements (Fig. **??**) indicates that the cell death rate is correlated to the number of baculovirus DNA copies in the cell with a logarithmic dependence. Rep78 concentration (*k*_*death,rep*_ = 0, Table 1) does not appear to significantly affect viability, in contrast to what has been found in mammalian cells.^50^ The cytolytic effect of rAAV capsids is also found to be negligible (*k*_*death,cap*_ = 0). The result that neither *Rep* nor *Cap* expression contributes significantly to the death rate of infected Sf9 cells is consistent with the literature finding that, following passage amplification, the titers of repBV, capBV and goiBV are similar.^52^

### Model Limitations

The model accurately reproduces several experimental datasets from the literature. However, the following limitations must be considered when using the model. Substrate and metabolite limitations due to depletion are not taken into account and neither are the effects of noxious metabolic products, under the assumption that the cellular macromolecular synthesis is shut-off soon after the process onset. The effect of metabolites on reaction kinetics is also not included in the model. Although results show that vector genome encapsidation is not limiting for the considered implementations of the BEVS, encapsidation could become limiting with different arrangements of the baculovirus construct. For simulating these scenarios with the model, a more accurate estimation of packaging kinetics should be conducted, improving the estimation of Rep52 limitation and, potentially, considering ATP limitation to encapsidation.^53^ While we validated the model for data coming from different laboratories, lab-to-lab variability is expected to affect the accuracy of the model predictions. Re-fit of model parameters, especially of viability-related parameters (*k*_*death,DNA*_ and *τ*_*death*_), would be needed to produce the most accurate predictions in a specific setup. Additionally, the parameters of the model have been estimated for the production of rAAV-2. While the model is able to track production dynamics for other serotypes (e.g., rAAV-5 in Fig. 5c), re-estimation of the parameters associated with vector genome amplification and encapsidation would be needed to improve the accuracy of model predictions for other serotypes.

### rAAV Production Modeling: Insect Cells vs. Mammalian Cells

Triple transient transfection (here referred to as *triple transfection*) of plasmid DNA into mammalian cells (especially human embryonic kidney 293 cells, HEK293) is the most common alternative to rAAV manufacturing via the BEVS.^6^ The reaction-transport network of processes based on mammalian and insect cells share several steps,^26^ but also have many differences that should be accounted for during process modeling.

First and foremost, insect cell-based processes involve an infection, which halts cell growth and leads to cell death within a few days. In contrast, cell cycle is not disrupted following transient transfection, with only modest loss of viability in suitable transfection conditions (plasmid concentrations, reagents, etc). This difference is probably the reason why Rep78 cytotoxicity affects viability of mammalian cells^50^ more than of baculovirus-infected insect cells, as we estimate. Other differences between infection and transient transfection processes involve DNA replication and infection propagation. Baculovirus DNA replicates to high levels upon infection, whereas the plasmid DNA that enters cells during transient transfection does not replicate, potentially providing less templates available for transcription. At the same time, baculovirus infection can self-propagate to uninfected cells through budding, while subsequent transient transfection waves can only occur upon introduction of fresh plasmid DNA into the culture medium.

The genes in the plasmids used for transient transfection and the genes introduced in recombinant baculoviruses in the BEVS have inherent differences. While a *Rep, Cap*, and ITR/GOI cassettes are present in the construct in both cases, transient transfection of mammalian cells requires the introduction of a helper plasmid, carrying genes of a helper virus that is needed for rAAV production. In the BEVS, instead, the baculovirus itself serves as helper virus, avoiding the need to introduce in the host additional helper genes. Furthermore, only *Rep52* and *Rep78* are expressed during rAAV manufacturing in the BEVS, whereas the spliced proteins Rep40 and Rep68 are also produced in transient transfection. Genes that lead to rAAV production are expressed in the BEVS through the baculovirus promoters, while in transient transfection the promoters of wild-type AAV are conventionally used. Finally, empty and filled capsids secrete into the cytosol and into the extracellular matrix in mammalian cells,^54^ facilitated by specific egress factors.^55^ Instead, no significant secretion is registered in insect cells, especially for empty capsids.^56^

Mammalian and insect cell processes have different efficiencies for rAAV production. In transient transfection, the typical filled-to-empty particles ratio is 5–30%.^7,8^ Instead, in the BEVS, filled particle ratios as high as 50–80% have been achieved.^10,19,48^ Recently, a comprehensive mechanistic model for rAAV manufacturing via transient transfection of HEK293 cells has been presented in the literature.^26^ The model indicated that a temporal lag between capsid synthesis and vector genome amplification plays a main role into the low efficiency of transient transfection in mammalian cells. This lag does not occur in insect cell processes (Fig. 7a), potentially explaining the difference in efficiency between the two systems. However, given the many differences between mammalian and insect cells processes, a more thorough differential model-based analysis should be conducted to understand which are the most critical steps that originate the efficiency gap. The results of the analysis can be leveraged to optimize the efficency of both systems.

## Conclusions

The current interest in achieving large production of rAAV viral vectors motivates the construction of a mechanistic model suitable for increasing understanding and to support process optimization. In this work, we develop the first mechanistic model for rAAV production in the BEVS. The model is successfully validated against experimental data from multiple key studies on rAAV manufacturing via the BEVS, including for TwoBac, ThreeBac, and OneBac constructs. We apply the model to thoroughly analyze production bottlenecks for TwoBac, one of the most widely used processes from the state-of-the-art. The model analysis indicates that larger rAAV production can be achieved by inducing stronger vector genome amplification. Stronger *Rep* expression, especially with respect to *Rep78*, can increase the rate of vector genome amplification, according to the model estimation. Incidentally, ThreeBac is found to be even more affected by Rep78 limitation to rAAV production than TwoBac. Reducing expression of the transgene and of baculovirus genes non-essential for the process can also lead to increased rAAV production, by diminishing the competition for *Rep* and *Cap* expression. Maintaining large cell viability for as long as possible during infection is crucial to achieve a high rAAV titer. The model-based analysis indicates that the cell death rate is correlated to the number of baculovirus DNA copies in the cell with a logarithmic dependence. Additionally, differently from mammalian cells, the viability dynamics of Sf9 cells expressing Rep78 is not different from the viability dynamics of other baculovirus-infected cells that do not express Rep78. The model indicates that vector genome encapsidation kinetics does not significantly limit rAAV production in TwoBac and ThreeBac. Finally, the model also suggests that a productive coinfection in which binding is not limiting production is achieved for MOI as low as 2 or 3 per each type of baculovirus. The model can be used to design and test in-silico different BEVSs with novel promoter-cassette arrangements, which can potentially lead to enhanced rAAV production.

## Materials and Methods

### Mathematical Model Formulation

#### Overview

For conciseness, the model formulation is first described in detail with reference to the TwoBac system. The ThreeBac and stable cell lines implementations of the model used for generating the results reported in this article are discussed in dedicated subsections below. The model equations reproduce the baculovirus infection dynamics and the reaction-transport network sketched in Fig. 1. The model is implemented in MATLAB (MathWorks, Waltham, MA, USA). In all simulations, the model equations are solved using the ordinary differential equation solver ode45 available in MATLAB. The model is developed for two-wave synchronous infection, and considers 9 viable extracellular species (Fig. 1b).

In addition to the concentrations in the culture of the repcapBV virion (*V*_*rc*_), the goiBV virion (*V*_*goi*_), and the viable uninfected cells (*T*), the model tracks the concentrations of viable infected and coinfected cells *I*_*j*_, considering cells infected by repcapBV (first wave: *j* = *rc*1, second wave: *j* = *rc*2), by goiBV (first wave: *j* = *goi*1, second wave: *j* = *goi*2), and of viable coinfected cells (first wave: *j* = *co*1, second wave: *j* = *co*2).

The model accounts for 13 intracellular species *X*_*j*_ for each type *j* of viable infected and coinfected cell. The intracellular species *X*_*j*_ are (Fig. 1c): receptor-bound repcapBV (*B*_*rc,j*_), receptor-bound goiBV (*B*_*goi,j*_), repcapBV DNA in nucleus (*vDNA*_*rc,j*_), goiBV DNA in nucleus (*vDNA*_*goi,j*_), *Rep* transcript (*mRNA*_*rep,j*_), *Cap* transcript (*mRNA*_*cap,j*_), transgene transcript (*mRNA*_*goi,j*_), Rep52 (*Rep*52_*j*_), Rep78 (*Rep*78_*j*_), non-encapsidated vector genome (*GOI*_*j*_), transgene protein (*GFP*_*j*_), rAAV empty capsids (*Caps*_*j*_), and rAAV filled capsids (*rAAV*_*j*_). The transgene protein is here denoted as GFP, since it is the most commonly used transgene in experiments for process development. The extension to any other transgene is straightforward. The concentration of all intracellular species is calculated for each infected and coinfected cell type, even though not all infections lead to production of the whole set of intracellular species (e.g., Rep proteins are produced only upon repcapBV infection). This implementation, although not optimal for speed, is designed to simulate more easily the different baculovirus constructs and experimental setups analyzed in this study (TwoBac, ThreeBac, and stably transfected cell lines).

Considering all viable extracellular species and the intracellular species in viable infected and coinfected cells, the model features a total of 87 species. Additional equations are set up for tracking the concentration of nonviable cells and of the respective intracellular species content. The intracellular species *X*_*j*_ is robustly expressed in the model as the cumulative concentration in the system across a given class *j* (#/mL). The total concentration in the system of a given intracellular species *X* is calculated as

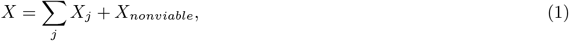

where *X*_*nonviable*_ is the cumulative concentration of the considered species in nonviable cells.

The phenomena considered by the model are: baculovirus binding, baculovirus transport to nucleus and replication, release of budded baculovirus, transcription and translation of AAV genes, rAAV capsids formation, Rep proteins synthesis, vector genome amplification, and vector genome encapsidation (Fig. 1c). Baculovirus infection presents an early (0–6 hpi), a late (6–18 hpi), and a very late (*>*18 hpi) stage, characterized by the activation of different reactions and mechanisms within the infected cell. For instance, phenomena such as DNA replication and budding occur only, respectively, in the late and very late stages (Fig. 1c). Since the model considers synchronous infection, the infection age of the first wave of infected and coinfected cells is set to the time passed from the process onset, whereas the infection age of the second wave is the time from the onset of budding for the first wave (*τ*_*rel*_).

For simplicity, the remainder of this discussion drops the two-wave notation, and the model is presented for a one-wave scenario, without any loss of generality. The notation for infected and coinfected cells *I*_*j*_ and for the concentration of the respective intracellular species *X*_*j*_ becomes *j ∈ {rc, goi, co}*. In the equations for nonviable cell concentration and the relevant content of intracellular species, not detailed for sake of conciseness, the only phenomenon that is accounted for is the degradation of intracellular species (with the same kinetics as in viable cells). Table 1 summarizes the model parameters, which are directly obtained from the literature or estimated from literature data, following the procedure outlined in the Parameter estimation strategy subsection.

#### Cells, virions, and binding

The balance for uninfected cells is

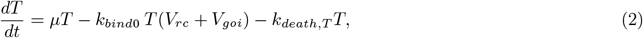

where *μ, k*_*bind*0_, and *k*_*death,T*_ are, respectively, the growth, binding, and death kinetic constants of infected cells. Infected cells do not undergo mitosis,^12^ and their balances are

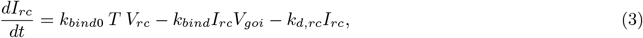

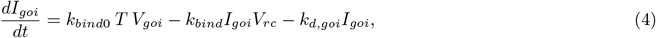

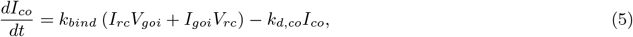

where *k*_*bind*_ is the equivalent binding kinetic constant for infected cells, and *k*_*death,j*_ (for *j ∈ {rc, goi, co}*) is the equivalent death kinetic constant of infected cells. We model the decay of binding rate that infected cells undergo with the infection progress as^12,57^

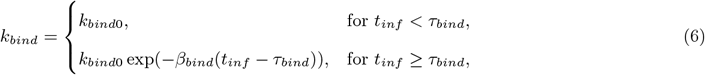

where *t*_*inf*_ is the infection age, *β*_*bind*_ is the binding decay kinetic constant, and *τ*_*bind*_ is the binding decay onset. The death rate of infected cells is faster than for uninfected cells, and most cells become nonviable between 3–5 dpi.^12,30^ The equivalent death kinetic constant of infected cells is modeled by

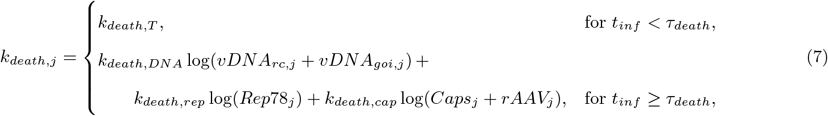

where *τ*_*death*_ is the infection age of switch from uninfected to infected cell death kinetics, and the parameters *k*_*death,DNA*_, *k*_*death,rep*_, and *k*_*death,cap*_ are kinetic constants relating *k*_*death,j*_ to intracellular concentrations of baculovirus DNA, Rep78, and rAAV capsids. Literature studies suggest that the concentration of these intracellular species might be correlated to the accelerated death rate of infected cells.^29,50^ However, following the parameter estimation findings reported in the Results section (specifically in subsection “Cell growth and death dynamics subsection”), the effect of *Rep* and *Cap* expression on *k*_*death,j*_ can actually be neglected (*k*_*death,rep*_ = *k*_*death,cap*_ = 0), and Eq. 7 can be simplified as

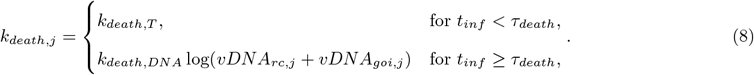

The balances for baculovirus virions are

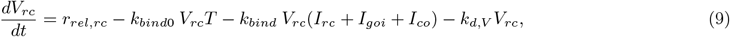

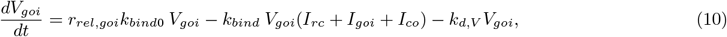

where *k*_*d,V*_ is the degradation kinetic constant for baculovirus, and *r*_*rel,rc*_ and *r*_*rel,goi*_ are the cumulative release rate of repcapBV and goiBV from infected cells, respectively. The release rates are given by

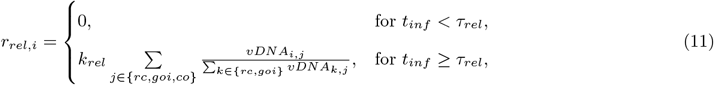

for *i ∈ {rc, goi}*, where *k*_*rel*_ is the release kinetic constant and *τ*_*rel*_ is budding onset time. The formulation of Eq. 11 assumes that infected cells release a fixed number of virions per unit of time during the very late stage and that, in case of coinfection, the virion production is split between repcapBV and goiBV, proportionally to the respective DNA copies in the cell. Although the literature approximately agrees on the values of *k*_*rel*_ = 9.8 *±* 1.5 PFU cell^*−*1^ h^*−*1^ and *τ*_*rel*_ = 18 *±* 2 hpi,^12,30^ different works suggest different end-points for budded virus release, ranging from *≈* 35 hpi^28^ up to cell death occurrence.^30^ For our model, we choose the latter option (Eq. 11). Since we only consider systems in which all cells are infected during the first or second synchronous infection waves, the budding end time does not practically affect the model simulation.

#### Transport to nucleus and replication

As for all intracellular species, the balances for the baculovirus bound to the receptors of viable infected cells are formulated in terms of cumulative concentration in the system:

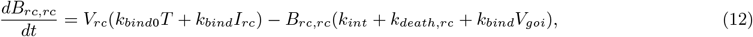

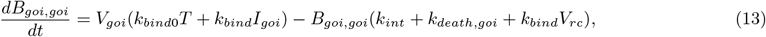

where *k*_*int*_ is the kinetic constant for internalization. Equations 12 and 13 include a source term, accounting for baculovirus binding, and a sink term, accounting for internalization, loss of viability, and for the amount of receptor-bound baculovirus that passes from the infected to the coinfected cells balance consequently to coinfection occurrence. The balances for the baculovirus bound to the receptors of viable coinfected cells present analogous contributions:

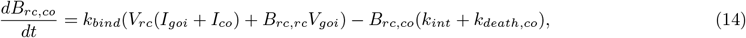

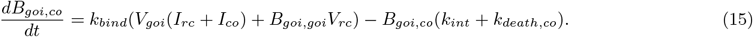

The balances for the viral DNA in the nucleus are

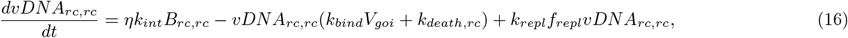

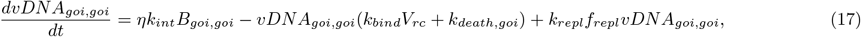

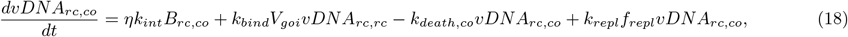

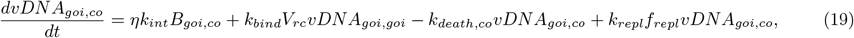

where *η* is a coefficient accounting for baculovirus degradation in lysosomes during trafficking to nucleus, *k*_*repl*_ is the baculovirus DNA replication kinetic constant, and *f*_*repl*_ is the activation function for baculovirus replication. Following the baculovirus infection dynamics:^12^

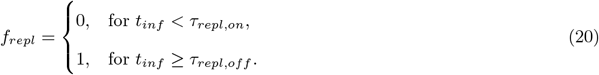

Beside DNA replication, the source contribution of Eqs. 16 to 19 accounts for internalization. As for receptor-bound baculovirus (Eqs. 12 to 15), Eqs. 16 to 19 present a contribution that factors out of the balance the baculovirus DNA in cells that become nonviable, and a contribution that moves the viral DNA content of infected cells that at a given time become coinfected from the infected cells balance to the coinfected cells balance. The latter contribution is necessary for closure, since binding, even though weakly, can still occur at the replication onset, and a cell infected by only one of the two baculoviruses might obtain coinfection status after having already started baculovirus replication. This contribution does not appear in the balances of the other intracellular species that are presented in the following discussion, since, when they form, baculovirus binding is weak or not happening anymore, preventing the flux of intracellular species from infected to coinfected cells balance. By definition, *B*_*goi,rc*_ = *B*_*rc,goi*_ = *vDNA*_*goi,rc*_ = *vDNA*_*rc,goi*_ = 0.

#### Transcription and translation

The model includes equations for expression of *Rep52, Rep78*, viral proteins, and the transgene. Transgene expression is explicitly considered, since it can potentially reduce the total host expression capability available for AAV structural and non-structural proteins. All the other proteins that are expressed during baculovirus infection are not featured in the model, as their expression pattern is not expected to affect rAAV production, nor to significantly vary across different experimental conditions.

The transcription of a generic gene *g* located in the DNA of a baculovirus of type *i* in the nucleus of infected cell of type *j* is modeled by

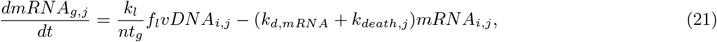

where *k*_*l*_ is the transcription kinetic constant for promoter *l* associated with gene *g, f*_*l*_ is the activation function for transcription from promoter *l, nt*_*g*_ is the number of nucleotides of gene *g*, and *k*_*d,mRNA*_ is the degradation kinetic constant for mRNA. As in the balances for receptor-bound baculovirus (Eqs. 16 to 19) and nuclear viral DNA (Eqs. 16 to 19), Eq. 21 accounts for the decrease of the total level in the system of mRNA available for translation due to loss of cell viability. For TwoBac, considering that *g ∈ {rep, cap, goi}, i ∈ {rc, goi}, j ∈ {rc, goi, co}*, and *l ∈ {polh, p*10*}*, and with the *g* : *i* : *j* : *l* mapping of Fig. 1a, Eq. 21 becomes

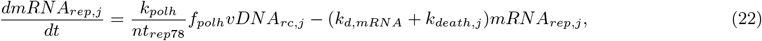

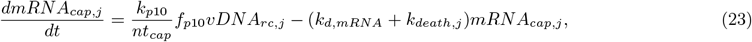

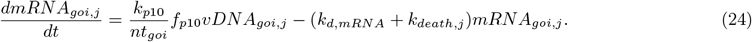

The activation function for the polh and p10 promoters are

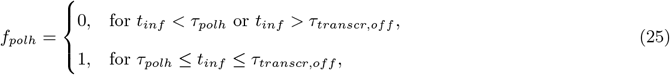

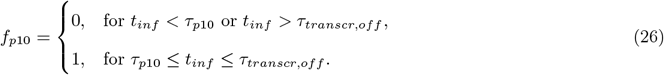

The translation rate *r*_*transl,g,j*_ for transcript *mRNA*_*g,j*_ in cells *j* is calculated, in terms of [# h^*−*1^ mL^*−*1^], with a Michaelis-Menten kinetics, accounting for saturation of the host translation machinery:

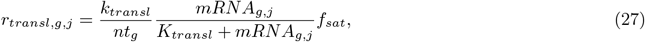

where *k*_*transl*_ is the translation kinetic constant, corresponding to the maximum translation rate, *K*_*transl*_ is the Michaelis-Menten constant for translation, and *f*_*sat*_ accounts for competition for translation among different transcripts:

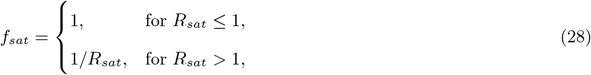

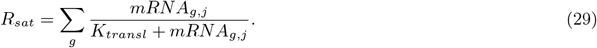

With the formulation of Eq. 27, translation spontaneously halts with mRNA degradation. As for transcription, for the TwoBac system, *g ∈ {rep, cap, goi}* and *j ∈ {rc, goi, co}* in Eqs. 27 to 29.

The corresponding balances for the Rep proteins are

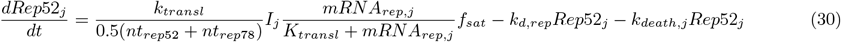

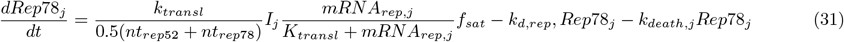

where *k*_*d,rep*_ is the degradation kinetic constant for Rep. In Eqs. 30 and 31, *nt*_*g*_ is taken as the average number of nucleotides between *Rep52* and *Rep78*, following the experimental finding that the leaky scanning mechanism of TwoBac leads to formation of Rep52 and Rep78 in ratio approximately equal to 1:1.^19^

The balance for the transgene protein is

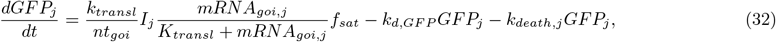

where *k*_*death,j*_ is the degradation kinetic constant for the protein.

Under the hypothesis of fast capsid assembly,^26,58,59^ the viral protein formation and assembly are lumped into a single step. The empty capsids balance becomes

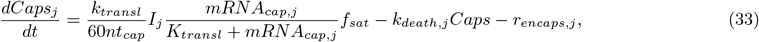

where *rencaps,j* is the vector genome encapsidation rate in infected cells of type *j*. This equation neglects capsid degradation, following experimental evidence from the literature,^42,44^ and considers that capsids protein VP1, VP2, and VP3 have the same relative synthesis rate, as assumed in previous models.^26^ Equation 33 does not account for empty capsid secretion into the cytosol, following experimental findings.^56^

#### Vector genome amplification and encapsidation

Rep78 plays a role the vector genome amplification process.^7,60^ Accordingly, vector genome amplification is modeled through a Michaelis-Menten kinetics, accounting for potential Rep78 limitation. The balance for the free vector genome available for encapsidation is

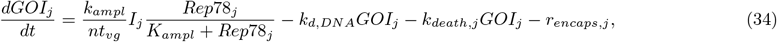

where *k*_*ampl*_ is the kinetic constant for amplification, *K*_*ampl*_ is the Michaelis-Menten constant, *nt*_*vg*_ is the number of nucleotides in the ITR/GOI cassette, and *k*_*d,DNA*_ is the degradation kinetic constant for the non-encapsidated vector genome. Equation 34 assumes that, in absence of Rep78 limitation, free vector genome is produced in the cell at the maximum rate of *k*_*ampl*_. Limitations induced by low levels of template are not considered, following experimental evidence.^47^

Vector genome encapsidation is mediated by Rep52, which forms stable intermediate complexes with the empty capsids, and intervenes in genome packaging.^7,60^ Rep52 limitation is modeled through a Michaelis-Menten kinetics, imposing the Michaelis-Menten constant in cells *j* (*K*_*encaps,j*_) equal to

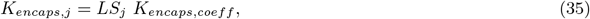

where *K*_*encaps,coeff*_ = 5, and *LS*_*j*_ is the limiting species for encapsidation in cells *j*:

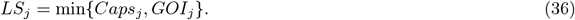

This modeling choice is based on the 1:1 mapping between Rep52 and capsids in the formation of intermediate complexes during encapsidation,^7^ assuming that Rep52 is not significantly limiting packaging when it is present in the cell in concentrations of one or more orders of magnitude larger than *LS*_*j*_. The resulting encapsidation rate is

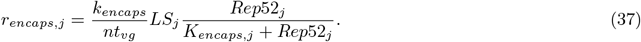

The corresponding balance for the filled rAAV capsids is

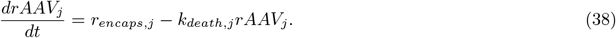

#### ThreeBac modeling

The baculovirus infection dynamics and the reaction-transport network for ThreeBac are shown in Fig. S1. ThreeBac modeling is carried out with the same rationale adopted for TwoBac, but accounting for the presence of a third baculovirus for gene delivery, and for the different arrangements and promoters of the *Rep, Cap*, and ITR/GOI cassettes (Fig. 1a). In ThreeBac, there are three virions (repBV, capBV, and goiBV), and 6 types of infected and coinfected cells *I*_*j*_ for each of the two synchronous waves (Fig. **??**). Balances for uninfected and infected cells and for the virions are analogous to the balances for TwoBac (Eqs. 2 to 11).

The ThreeBac model accounts for 16 intracellular species, which is three more than for TwoBac. Two additional intracellular species are related to the presence of one more type of receptor-bound baculovirus and baculovirus DNA in the nucleus, whereas the third additional species originates from the separate transcription of *Rep52* and *Rep78* in ThreeBac (Fig. **??**). Balances for receptor-bound baculovirus and baculovirus DNA in the nucleus are developed accordingly to the TwoBac model (Eqs. 12 to 20). The balances for the transcripts are derived from Eq. 21, accounting for the promoter-cassette arrangement of ThreeBac:

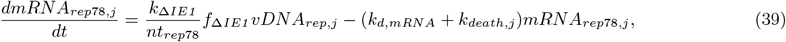

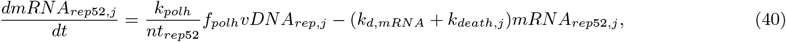

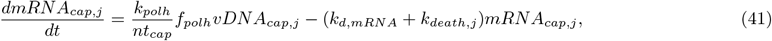

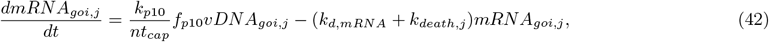

where *k*_Δ*IE1*_ is the transcription kinetic constant for the Δ*IE1* promoter, and *f*_Δ*IE1*_ is the promoter’s activation function:

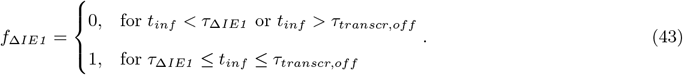

Translation, Rep, Cap and transgene protein synthesis, capsid formation, vector genome amplification, and encapsidation are modeled as in the equations for TwoBac (Eqs. 27 to 38).

#### Stably transfected cell lines modeling

The model considers two different configurations for rAAV production through baculovirus infection of stably transfected cell lines. The first configuration involves infection with repBV of a cell line with an integrated ITR/GOI cassette.^47^ This scenario is directly simulated using the same implementation of the model used for ThreeBac, imposing the concentration of goiBV DNA in the nucleus equal to one copy per cell, and the concentration of capBV and goiBV virions equal to zero. The second configuration involves the OneBac process, in which goiBV is used to infect a cell line stably transfected with *Rep* and *Cap* cassettes.^49^ The OneBac process is simulated using the model implementation for TwoBac, imposing the concentration of the repcapBV virion equal to zero. In the OneBac experiment considered in this work, weak *Rep* and *Cap* amplification is registered within the cell during baculovirus infection.^9^ Hence, in Eq. 21 the number of viral DNA templates per cell available for transcription is set equal to the *Rep* and *Cap* copy numbers measured during the experiment (respectively, about 50 and about 450 copies per cell).

### Parameter Estimation Strategy

The model has 32 parameters (Table 1). Parameter estimation is carried out through maximum likelihood estimation in the log space of the parameters:

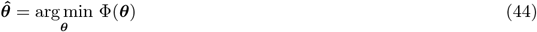

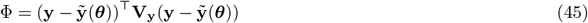

where ***θ*** is the vector of parameter estimates, **y** is the vector of measurements, 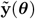 is the vector of model prediction for the measured variables, and *V*_**y**_ is the measurement covariance matrix. The covariance matrix of the estimated parameters (**V**_***θ***_) is computed through the Hessian approximation:^61^

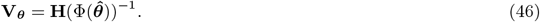

The parameters are estimated through a sequential approach, exploiting, at each step, the parameters that have been obtained at the previous step, as further outlined. Unless otherwise specified, in each step, a multi-start approach with a sequential quadratic algorithm is used to solve the optimization (Eqs. 44 and 45), by minimizing the objective function (Eq. 45) locally from 1,000 random starting points. The Hessian for computing **V**_***θ***_ (Eq. 45) is directly obtained from the output of the solver (MATLAB’s fmincon). First, we retrieve the parameters directly available in the literature, as reported in Table 1. Then, we estimate the binding parameters (*k*_*bind*0_, *β*_*bind*_, and *τ*_*bind*_) from the binding rate experimental measurements reported by Nielsen,^31^ and the baculovirus DNA replication parameters (*k*_*repl*_, *τ*_*repl,on*_, and *τ*_*repl,off*_) from the measurements of baculovirus DNA concentration during baculovirus infection reported by Rosinski et al.^38^ We estimate the parameters for transcription from polh (*k*_*polh*_ and *τ*_*polh*_) and p10 (*k*_*p*10_ and *τ*_*p*10_), the time of transcription halt (*τ*_*transcr,off*_) for all promoters and the mRNA degradation kinetic constant (*k*_*d,mRNA*_) from mRNA concentration and viability measurements collected in experiments for VLP production with the BEVS by Vieira et al.^39^ Upon analysis with ImageJ of western blot data reported in the literature,^19,41,42^ we set *k*_Δ*IE*1_ = *k*_*polh*_*/*10. We subsequently estimate the degradation kinetic constants for the non-encapsidated vector genome (*k*_*d,DNA*_) and for Rep proteins (*k*_*d,rep*_) from measurements of vector genome copy number^47^ and Rep concentration^42,46^ collected during the last stages of baculovirus infection. We finally estimate the remaining model parameters (*k*_*transl*_, *K*_*transl*_, *k*_*ampl*_, *K*_*ampl*_, and *k*_*encaps*_) from the measurements reported by Aucoin et al.^45^ of total capsids and rAAV at the end of nine batches carried out with the ThreeBac system. We carry out this parameter estimation twice. The first time, we use in the simulation different infected cell death kinetic parameters for every single batch, roughly estimated to match the viability measurements reported by Aucoin et al.^45^ Having now available the parameters for translation, amplification, and encapsidation, we can use the model to simulate the concentration of Rep78 and rAAV capsids during the experiments, which, in our model, affect the cell death dynamics (Eq. 7). We then regress a single set of parameters (reported in Table 1) for the infected cell death kinetics (*k*_*death,cap*_, *k*_*death,DNA*_, *k*_*death,rep*_, and *τ*_*death*_) through maximum likelihood estimation, from the viability measurements in the nine batches in the study reported by Aucoin et al. We then re-estimate the kinetic parameters for translation, vector genome amplification and encapsidation (*k*_*transl*_, *K*_*transl*_, *k*_*ampl*_, *K*_*ampl*_, and *k*_*encaps*_) on the same dataset, but using the (same) updated death kinetic parameters in all nine batches. This time, we carry out parameter estimation through bootstrap^62^ (500 samplings). The obtained parameters are reported in Table 1, and do not present a large difference compared to the parameters estimated during the first round.

The set of 32 parameters obtained through parameter estimation are used in the simulations for model validation (Figs. 2b, 3, 5, and **??**) and for in-silico analysis of the process (Figs. 6 and 7). The only exception are the parameters for the infected cell death kinetics (*k*_*death,DNA*_ and *τ*_*death*_), which, for model validation, are re-estimated to match the viability measurements in each experiment, when available. The resulting parameters (not reported) present a maximum deviation of 50% from the values reported in Table 1, indicating that viability can be much affected by lab-to-lab variability.

The sensitivity 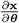 of the model states **x** with respect to the model parameters ***θ*** is computed by integrating the sensitivity equations alongside the model equations:^63^

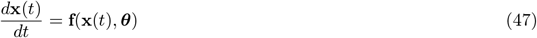

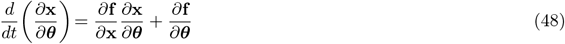

The Jacobians 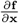 and 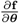 are calculated using the automatic differentiation toolbox ADiGator.^64^ The normalized cumulative sensitivity *S*_*x,θ*_ across the batch of a state *x* to a parameter *θ* is computed from

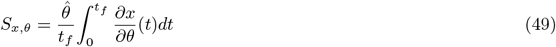

where *t*_*f*_ is the batch duration. The uncertainty of the model prediction is computed with a forward Monte Carlo approach with 5 *×* 10^4^ realizations, sampling, for each realization, a different set of model parameters from the distribution defined by the parameter estimate and variance matrix (Table 1).

Data are extracted from the figures of the cited literature using WebPlotDigitizer or ImageJ.

## Availability of Code

A MATLAB implementation of the model presented in this manuscript can be downloaded at the following link: https://github.com/francescodestro/rAAV_BEVS.

## Acknowledgements

This work was done in Cambridge, MA, USA. This work was supported by the US Food and Drug Administration through Contract no. 75F40121C00131.

