## Supplementary Material for "Mechanistic Modeling Explains the Production Dynamics of Recombinant Adeno-Associated Virus with the Baculovirus Expression Vector System"

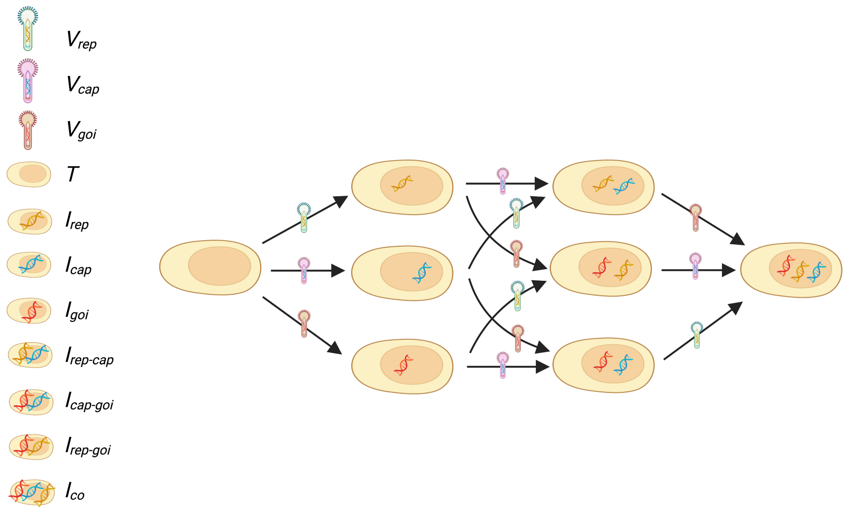


**(a)**

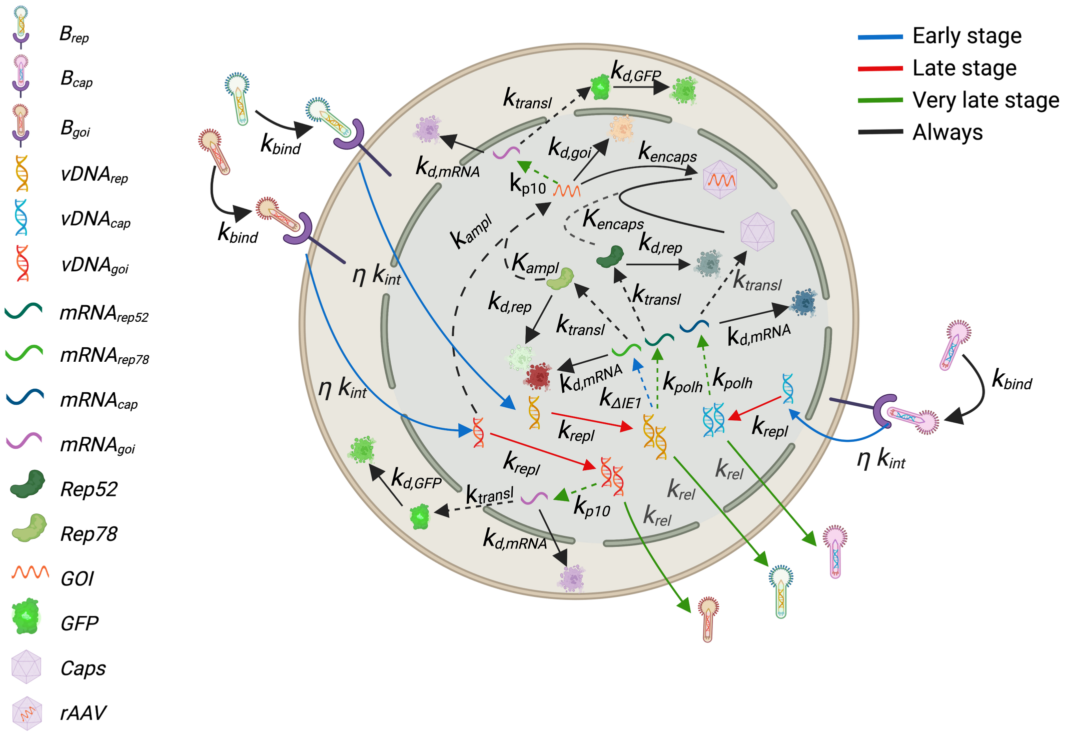


**(b)**

**Figure S1:** ThreeBac: (a) Baculovirus infection dynamics: coinfection by all three baculoviruses is necessary for rAAV production. (b) Intracellular reaction-transport network. Differently from TwoBac, *Rep* and *Cap* are delivered through two separate baculoviruses, and Rep proteins are expressed through separate *Rep52* and *Rep78*. Dashed lines indicate that reactants are not consumed in the reaction. Figures were created with BioRender (https://biorender.com/).

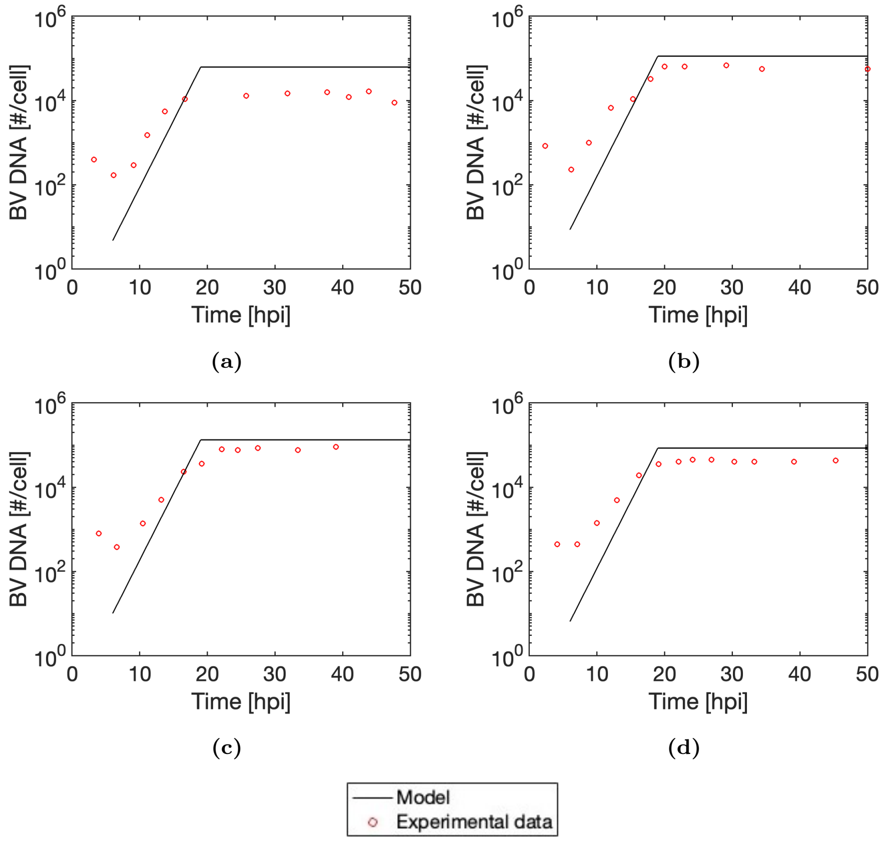


**Figure S2:** Model fit to baculovirus DNA copy number per cell during baculovirus infection from the dataset[35] used for estimation of $k_{repl}$. Experiments are carried out at MOI=20 and cell density at the time of infection equal to (a) $5 \times 10^5$ cell mL$^{-1}$ (b) $1 \times 10^6$ cell mL$^{-1}$ (c) $3 \times 10^6$ cell mL$^{-1}$ and (d) $5 \times 10^6$ cell mL$^{-1}$.

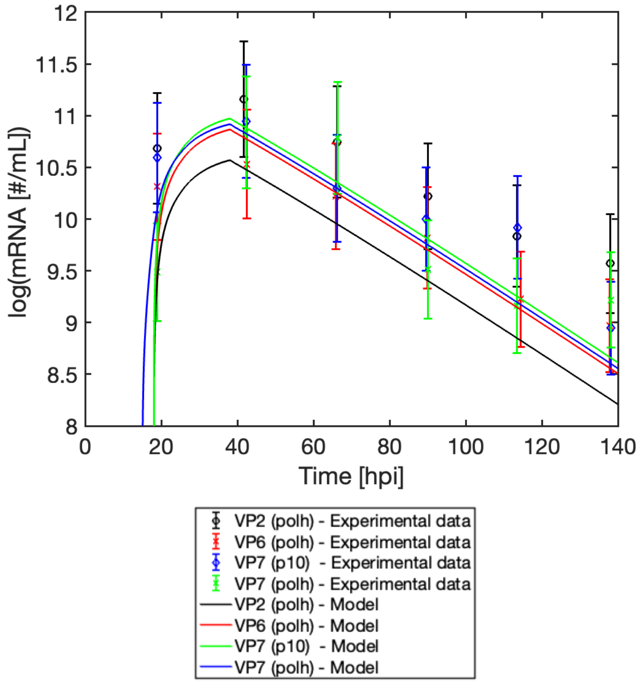


**Figure S3:** Model fit to transcript concentration of rotavirus structural proteins VP2, VP6 and VP7 during experiments of rotavirus virus-like-particle manufacturing with the BEVS from the dataset[39] used for estimation of $k_{polh}$, $k_{p10}$, $\tau_{polh}$, $\tau_{p10}$ and $k_{d,RNA}$. Experimental concentrations for VP2 and VP6 are reported as the average between the two experimental runs contained in the dataset.

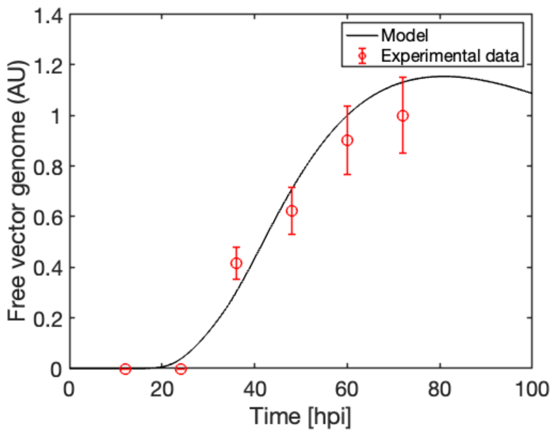


**Figure S4:** Model validation: free vector genome available for encapsidation. Data from Figure 4c (ΔIE) in Urabe et al.[41]

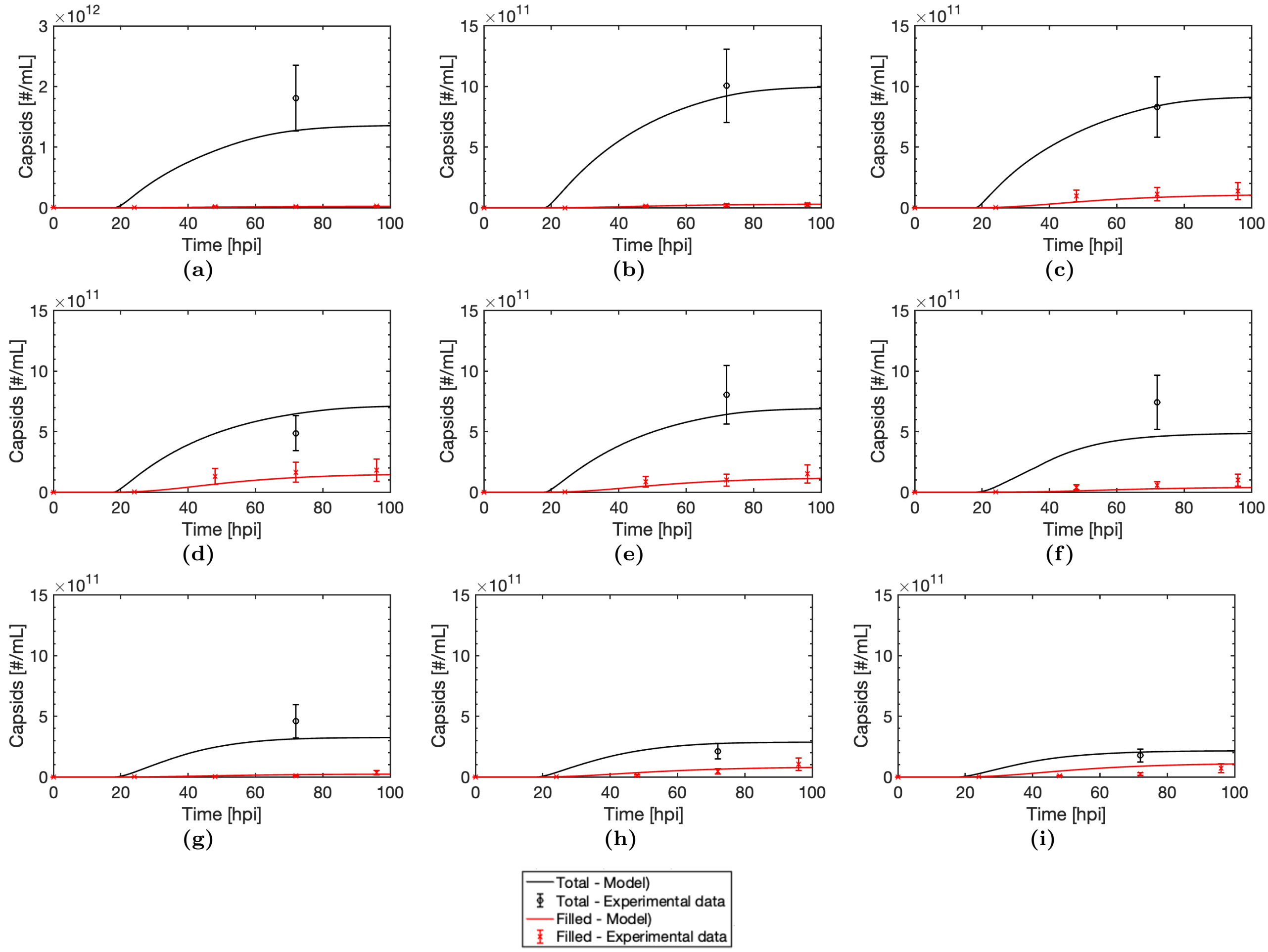


**Figure S5:** Model fit to ThreeBac dataset[44] used for estimation of $k_{transl}$, $K_{transl}$, $k_{ampl}$, $K_{ampl}$ and $k_{encaps}$: total capsids at 72 hpi and dynamic measurements of rAAV filled capsids concentration across the batches. Only the measurements of total capsids at 72 hpi and of rAAV filled capsids at 96 hpi are used for parameter estimation. The reported filled capsids concentration measurements are obtained by multiplying the infective viral particle titer reported in Figure 2a in Aucoin et al.[44] by a factor of 1000. The dataset includes nine batches, in which the MOIs of capBV, goiBV and repBV are, respectively: (a) 9, 1, 1 (b) 9, 9, 1 (c) 9, 1, 9 (d) 9, 9, 9 (e) 5, 5, 5 (f) 1, 1, 1 (g) 1, 9, 1 (h) 1, 1, 9 (i) 1, 9, 9.

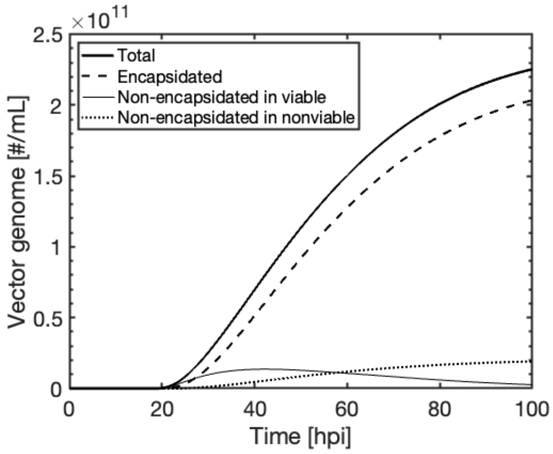


**Figure S6:** In-silico analysis of the process: vector genome encapsidation analysis for TwoBac. Comparison of total number of replicated vector genome vs encapsidated vector genome vs non-encapsidated vector genome in viable cells vs non-encapsidated vector genome in nonviable cells.

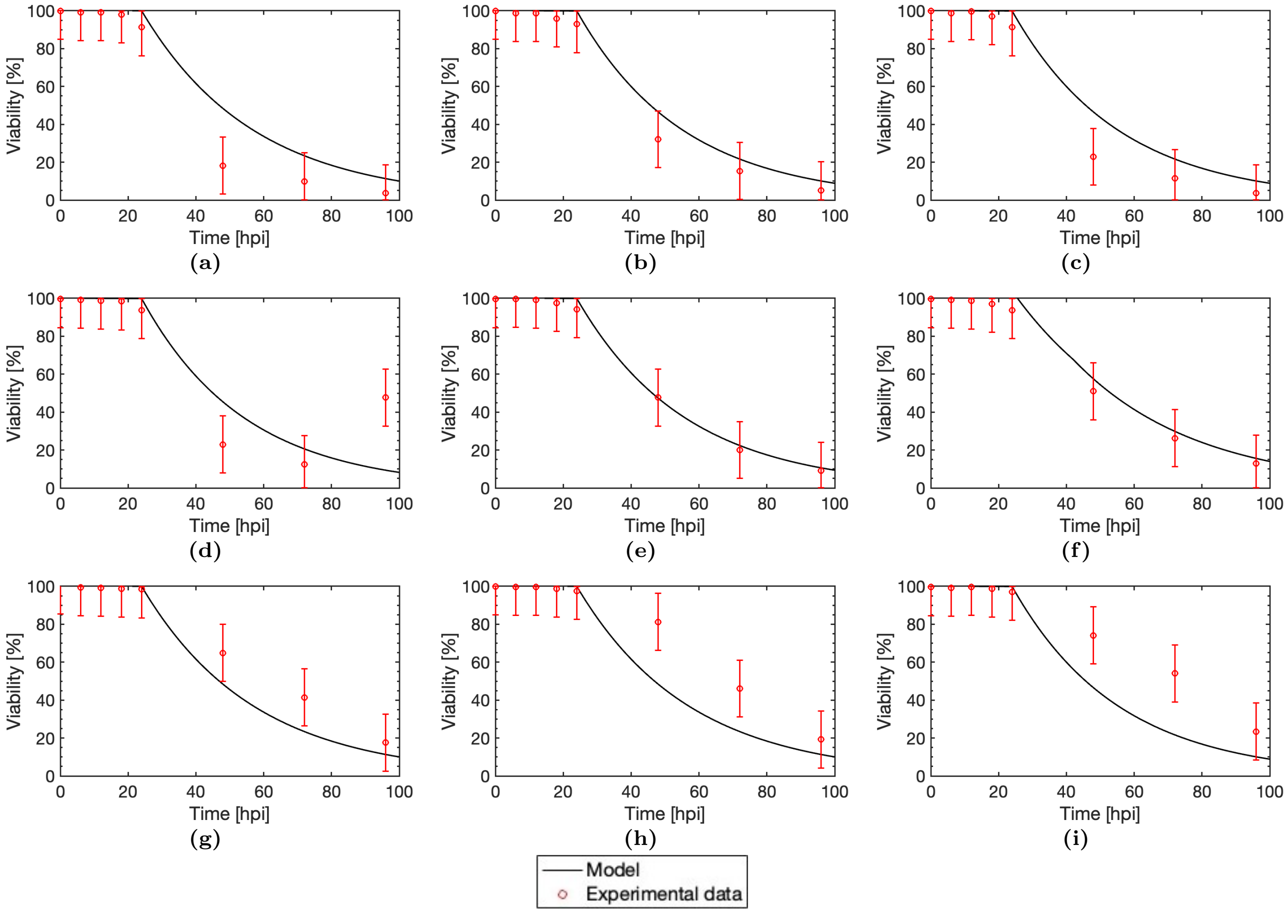


**Figure S7:** Model fit to viability measurements from ThreeBac dataset[44] used for estimation of $k_{death,cap}$, $k_{death,DNA}$, $k_{death,rep}$ and $\tau_{death}$. The dataset includes the same nine batches considered in Fig. S5, in which the MOIs of capBV, goiBV and repBV are, respectively: (a) 9, 1, 1 (b) 9, 9, 1 (c) 9, 1, 9 (d) 9, 9, 9 (e) 5, 5, 5 (f) 1, 1, 1 (g) 1, 9, 1 (h) 1, 1, 9 (i) 1, 9, 9.
